# The genomic landscape of western South America: Andes, Amazonia and Pacific Coast

**DOI:** 10.1101/505628

**Authors:** Chiara Barbieri, Rodrigo Barquera, Leonardo Arias, José R. Sandoval, Oscar Acosta, Camilo Zurita, Abraham Aguilar-Campos, Ana M. Tito-Álvarez, Ricardo Serrano-Osuna, Russell Gray, Paul Heggarty, Kentaro K. Shimizu, Ricardo Fujita, Mark Stoneking, Irina Pugach, Lars Fehren-Schmitz

## Abstract

Studies of Native South American genetic diversity have helped to shed light on the peopling and differentiation of the continent, but available data are sparse for the major ecogeographic domains. These include the Pacific Coast, a potential early migration route; the Andes, home to the most expansive complex societies and to one of the most spoken indigenous language families of the continent (Quechua); and Amazonia, with its understudied population structure and rich cultural diversity. Here we explore the genetic structure of 177 individuals from these three domains, genotyped with the Affymetrix Human Origins array. We infer multiple sources of ancestry within the Native American ancestry component; one with clear predominance on the Coast and in the Andes, and at least two distinct substrates in neighboring Amazonia, with a previously undetected ancestry characteristic of northern Ecuador and Colombia. Amazonian populations are also involved in recent gene-flow with each other and across ecogeographic domains, which does not accord with the traditional view of small, isolated groups. Long distance genetic connections between speakers of the same language family suggest that languages had spread not by cultural contact alone. Finally, Native American populations admixed with post-Columbian European and African sources at different times, with few cases of prolonged isolation. With our results we emphasize the importance of including under-studied regions of the continent in high-resolution genetic studies, and we illustrate the potential of SNP chip arrays for informative regional scale analysis.

## INTRODUCTION

The genetic diversity of the Americas has long been underestimated due to the paucity of population samples analyzed with high resolution markers. Over the past two decades, population studies have focused on uniparental markers, predominantly typed at low resolution (reviewed in ^1^). Recent studies are increasing the coverage of the continent with high resolution genomic data from ancient remains and living populations. The results confirm previous finding at a continental scale, such as a post-Last Glacial Maximum entry of a small founding population, a major migration ancestral to all living Native American groups from North to South America ^2–6^, with further layers of population structure and admixture suggested by the analysis of ancient DNA. These demographic dynamics include an early diverging branch reconstructed for ancient North American sites ^7^, which did not reach South America ^8^, and an enigmatic signal of Australasian ancestry recovered only in some populations of South America ^9,10^. The early population differentiation experienced after the initial entry in the continent resulted in different ancestries, such as the “Mixe” ^10^ or the “ancient Californian Channel Islands” ^8^, as reconstructed by admixture graph methods. It is difficult to trace how these ancestral genetic components have survived in living populations, as there is a lack of dense sampling of populations with a high proportion of Native American ancestry. This also impacts our understanding of pre-colonial local scale dynamics, with only a few studies reporting a good sampling coverage for targeted regions ^11,12^.

In South America, genetic studies robustly recovered a substantial differentiation between the Andes and Amazonia, which has been framed within a model of large communities connected by gene-flow in the Andes vs. small, isolated communities in Amazonia ^13–15^. This model builds on evidence for major complex societies in the Andes (culminating with the well-known but short-lived Inca empire) which fostered population movements and connections, counterbalanced by the traditional view of the Amazon basin as the homeland of small, isolated tribes. The latter view is challenged by increasing evidence of large-scale societies ^16,17^, the role of rivers as primary routes for gene-flow ^12^, and the presence of important centers of plant domestication ^18^. To gain a better representation of the highly diverse cultural landscape of Amazonia, a more intense archaeological effort is needed, together with a more fine-grained sampling of living and ancient human populations.

In particular, this model of South American genetic structure typically overlooks the Pacific Coast, a key context for the early migration history of the continent ^19^ and the cradle of the earliest complex societies in South America, from 3000 BCE ^20^. Recent studies have begun to investigate human variation on the Pacific coast through aDNA ^21–23^ and by sampling urban areas ^11,24^, but to fill out this picture requires further, complementary genetic studies on living populations (especially from non-urban areas).

Language diversity can also be a factor shaping population relatedness. The diffusion of major language families is traditionally associated with demographic movements ^25,26^: this association was validated with genetic data for some of the largest language families of the world ^27–29^, but no strong candidates are found in South America, where genetic diversity overall does not correlated with linguistic diversity ^30^. Previous genetic work ^31,32^ evaluated alternative models of cultural vs. demographic diffusion for Quechua, the most spoken language family of the Andes, present also in small pockets of the Amazonian lowlands ^33^. These studies, based on uniparental markers, revealed intense contact routes in the southern highlands, but not in the northern regions nor in neighboring Amazonia. Relatively few genetic studies have addressed the diffusion of the main language families of Amazonia (notably Arawak, Tupí or Carib), although very recent research does focus on sub-branches or smaller families ^12,34^. Some scholars suggest a major role for cultural contact alone behind the diffusion of Arawak ^35^. The particularly fragmented distribution of the three major language families across much of lowland South America ^36^ calls for a more fine-grained sampling to test for potential connections between their speaker populations.

Here we focus on western South America to investigate environmental and cultural influences on population genetic structure over the three ecogeographic domains: the Andes, the Amazonia and the Pacific Coast, and transitional environments in between. The analysis of new genetic samples from populations with different cultural, linguistic and historical backgrounds should contribute to understanding both the modes of early migration and the immediate consequences of colonial contact. A first open question concerns the scale of the genetic impact of major complex societies, which arose in two main focal regions: the north coast of Peru, and the Andean highlands of Central and Southern Peru and northern Bolivia. Large-scale societies possibly left a trace in the demographic profiles of indigenous populations, associated with high population density ^37^, but the extent of long-distance population movements and the origins of the populations that developed such societies remain largely unknown. A second open question concerns the diffusion mechanisms of major language families. We aim at tracing genetic connections over the scattered and widespread diffusion of representative Andean and Amazonian languages, focusing in particular on a vast region where different varieties of Quechua are spoken. Finally, a third open question concerns the demographic events occurring over the last five centuries since European contact, and how they impacted upon different South American populations. Gene flow from European and African sources can be easily distinguished in the genomic ancestry of the American populations ^38,39^. The timing and intensity of the European-mediated admixture has been estimated for urban, heavily admixed regions ^11,40^, but has yet to be investigated systematically across South America.

## RESULTS

### Continental-scale population structure

Continental ancestry structure was investigated by means of ADMIXTURE analysis. Yoruba and Spanish population samples were included to distinguish admixture from European and African sources (Fig. 1, Fig. S2). The most supported value of K was 3 (Fig. S2), indicating that the clearest ancestry signal is the one that separates African, European and a shared Native American ancestry. Further levels of K were considered to explore structure within the Native American component. At K=4 a new component is found in most Amazonian populations, while at K=5 the Xavante are distinguished from the other populations (consistent with their high levels of genetic drift, as described by the diversity values discussed later). At K=6 the Amazonian populations divide further into one component common to the Kichwa from northern Ecuador (Kichwa Orellana) and the neighboring Colombian populations from the eastern slopes of the northern Andes (the light green ancestry component in Fig. S2, designated “Amazonia North”) distinct from a further component common to the remaining Amazonian populations (dark green ancestry component, designated “Amazonia Core”). At K=7 a further ancestry component is distinguished in the Central-Southern Andes (dark blue). At K=8 the North American populations are distinguished by a separate component (purple color), and at K=9 a component appears in the Colombian populations from the north-west (Kamentsa, Inga and Cofán, in light yellow), separating them from Kichwa Orellana, although at this level of K the cross-validation error begins to increase appreciably.

**Fig. 1.**
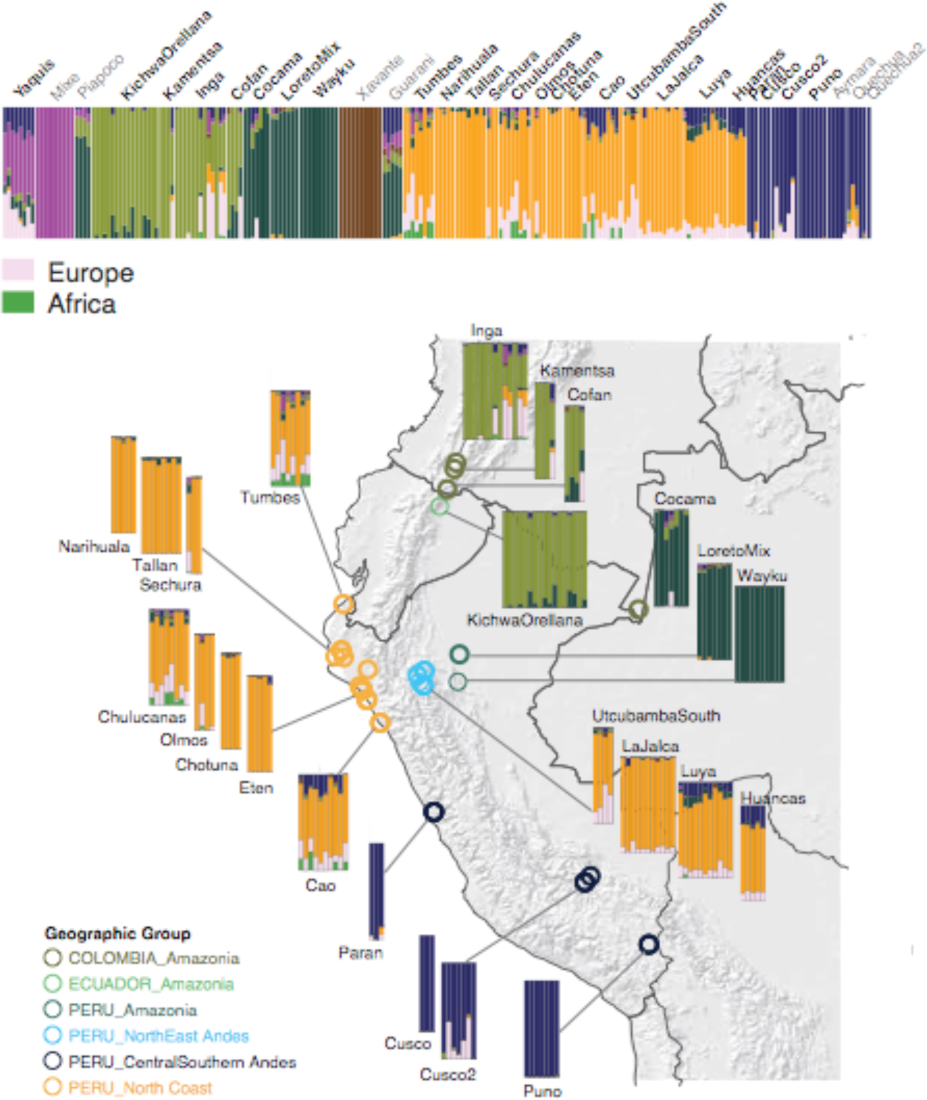
Map showing the approximate sampling locations of the newly reported population samples from South America, together with the ADMIXTURE results for K= 7. On top of the Admixture plot, newly reported population samples are in boldface.

This structure is reproduced in the PCA analysis, performed with the full set of SNPs as well as with a set of SNPs ascertained for Karitiana in the initial Human Origins assembly — Panel 7 in ^41^ — as shown in Fig. 2 and Fig. S3. In the full set PCA (Fig. 2) the first dimension is driven by European admixture, which pulls individuals with the highest amount of non-Native American component off to the right side of the plot. The second dimension already distinguishes the two Amazonian components, i.e. “Amazonia Core” and “Amazonia North”. The third dimension in the full set separates off individuals from the North Coast who have African admixture (Fig. S3A). The PCA with the ascertained set (Fig. S3B) is less influenced by the European and African signal, and is able to illustrate how the Native American structure coincides with geographical macro-areas. The first dimension separates samples from both “Amazonia North” and “Amazonia Core” from the rest of the Americas. The second dimension separates off “Amazonia Core”, the third separates the North American samples, and the fourth dimension separates the Central-Southern Andes. Individuals from different locations on the North Coast display a wide range of variation and in all dimensions partially overlap with the North-East Andes of Peru.

**Fig. 2.**
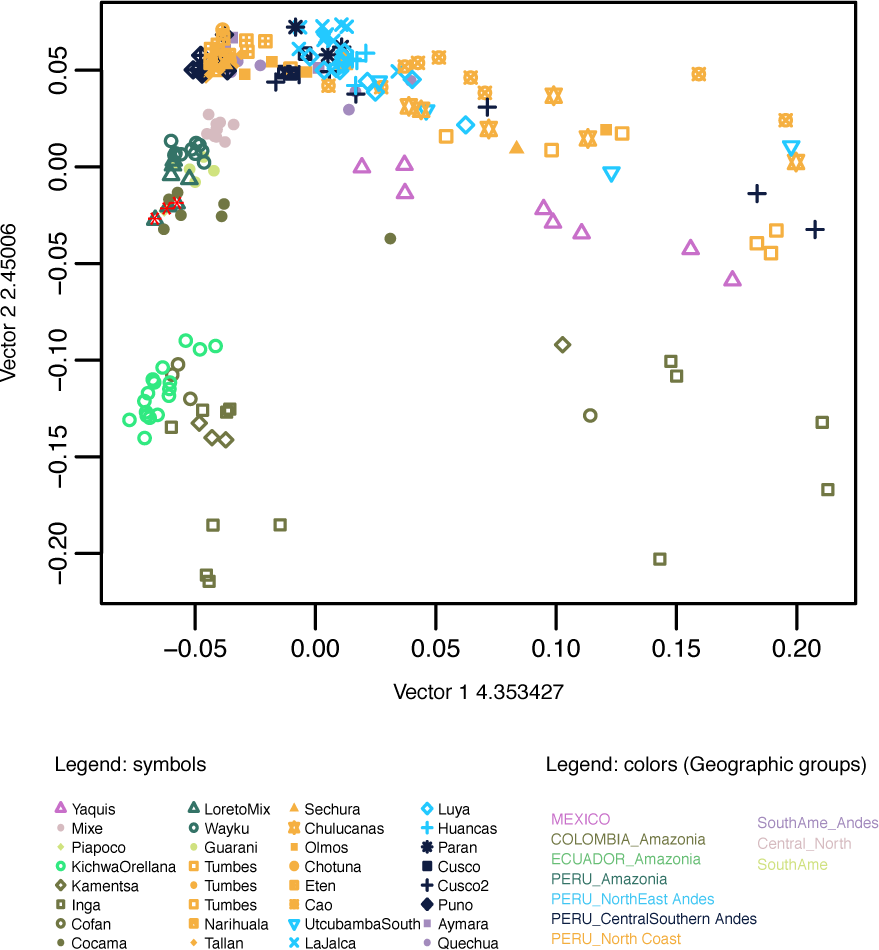
Principal Component Analysis of the newly reported samples together with representative populations from North and South America. Color legend corresponds to geographic grouping. Three Cocama-speaking individuals from the “LoretoMix” population are marked with a red asterisk in the first PCA panel and discussed in the section on IBD analysis.

Broad population relationships can be estimated by the F_ST_ distances between populations, here visualized with a NJ tree (Fig. S4). Outliers such as Chukchi, Pima, Cabecar, Xavante and Karitiana (which were excluded from the PCA analysis but are included here) exhibit very long and diverging branches. Populations from the Coast and the two regions of the Andes clearly belong to the same branch, but cluster separately. Yaquis and Pima (which both speak Uto-Aztecan languages) cluster together, separated from other populations from southern North America such as Mixe and Cabecar. Among Amazonian populations, Inga, Kamentsa, Cofán and Kichwa Orellana form one branch, while the other Amazonian populations also group together. The population structure therefore corresponds to a broad division between the following macro-regions: North America, Pacific Coast + Andes, and Amazonia, the last of which can be further divided between the proposed “Amazonia North” and “Amazonia Core” components.

### Demographic reconstructions and drift

To assess if we can distinguish different demographic trajectories for the populations considered, we analyzed Runs of Homozygosity (ROH) blocks. ROH blocks come from a shared ancestor, with their length inversely proportional to the number of generations since the split. ROH blocks that result from a recent bottleneck will tend to be longer, as there are fewer recombination events. ROH blocks are also informative about effective population size (N_e_), as populations with low N_e_ tend to have more extended ROH than those with high N_e_ ^42^. Here ROH blocks were divided either into two length classes as suggested by ^43^ (Fig. 3), or six bins as in a previous study of Native American populations ^34^ (Fig. S5). All of the populations have an excess of short ROH (<1.6 Mb); this excess was lower in those populations exhibiting more European admixture (Fig. 3). The short ROH likely reflect the strong bottleneck experienced by the founding population of the Americas, as previously noted ^34,43,44^. The long ROH classes are differently distributed among populations, regardless of their geographic proximity or, more broadly, their ecogeographic domain. The populations with the highest proportion of large ROH are the Karitiana, Xavante, Cabecar and Pima. We can further distinguish populations with fewer ROH blocks longer than 2-4 Mb (Group 1 in Fig. S5). Some of these populations have more European/African admixture (group 1a), but more interesting are the populations that are less admixed with Europeans (groups 1b and 1c in particular). The absence of long ROH implies that these populations did not share a recent bottleneck: in this group are populations from Amazonia (Cocama), most of the populations from the Coast, some from the Andes (La Jalca, Cusco2, Parán, Puno) and the Yaquis from Mexico. Populations with a peak of ROH length at 4-8 Mb may have experienced a recent bottleneck (Group 2): this is common in Amazonia (Kichwa Orellana from Ecuador, Cofán in Colombia, LoretoMix in Peru). Finally, a few populations from all three ecogeographic domains show a low peak at the longest ROH, 8-16 Mb (Group 3): this could indicate an even more recent population size reduction and/or isolation (See Fig. S5 for all of the individual population profiles per group).

**Fig. 3.**
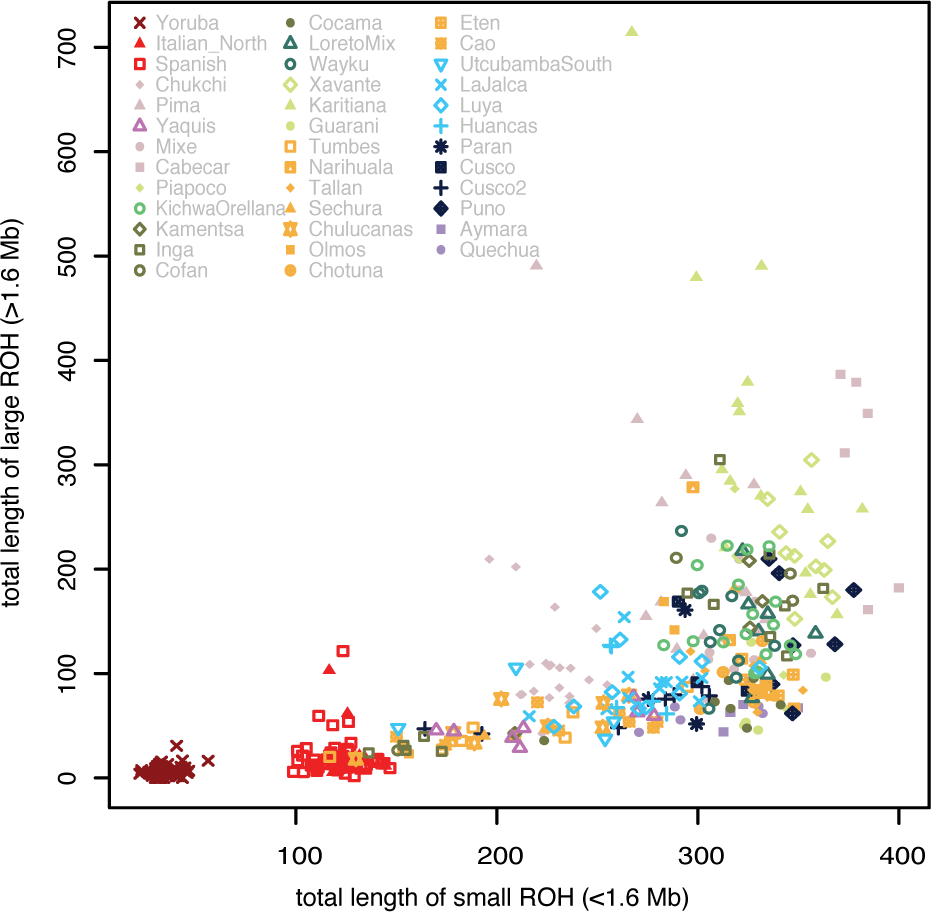
Proportion of small and large ROH classes for each individual.

Population internal diversity and drift was also evaluated by calculating estimates of consanguinity per individual (coefficient F), visualized in Fig. S6A. Published data for the Karitiana, Xavante and Cabecar have the highest levels of consanguinity. Overall, consanguinity is slightly higher in Amazonian populations than on the Coast. It is lower in the Andes, and lower in populations more admixed with non-American sources, such as North-East Peruvian Andes and the admixed populations of the Coast. The Puno population has the highest F values and the most ROH blocks of the Andean populations. The level of consanguinity appears to be directly correlated with the proportion of Native American ancestry (as estimated by Supervised ADMIXTURE analysis): Fig. S6B displays this correlation, with (as expected) slightly lower values of F by proportion of Native American ancestry on the Coast and in the Andes, and slightly higher F values in Amazonia, with a few individuals from the Inga, Yaquis, and Cusco2 populations who have less Native American ancestry but high levels of consanguinity.

### Recent contact from haplotype sharing networks mirrors linguistic connections

To investigate recent historical layers of contact we analyzed Identity by Descent (IBD) segments. Identical blocks between individuals correspond to shared ancestry, with longer blocks corresponding to recent shared ancestry. Fragments shorter than 5 cM are shared between almost all pairs of Native American populations (data not shown), in agreement with other studies of South American populations ^11^. This diffused pattern of sharing might reflect the reduced genetic diversity of the continent from the initial founding bottleneck (resulting in a high overall level of consanguinity, see ^44,45^). To focus on the most relevant sharing patterns, a threshold of 5cM was applied, and population pairs which shared only one fragment were not considered.

Fig. 4A shows the overall pairwise sharing patterns, while Fig. 4B includes only those pairs that have a number of shared blocks (adjusted for population size) higher than the median, to further highlight the most significant sharing networks. The highest impact of sharing events can be found along the diagonal in Fig. 4A, within the various regions covered: Amazonia, North Coast, North-East Andes and Central-Southern Andes. The network within the North-East Andes extends over a limited geographic distance between villages in the Chachapoyas province (<80km distant from each other), while the networks within the Coast and within the Central-Southern Andes both involve more long-distance sharing. Sampling locations along the Coast, where the total length of shared blocks is greater, span a longitudinal distance of almost 700 km, while sampling locations in Central-Southern Andes, where the total length of shared blocks is lower, cover a total distance of ∼1000 km. We find a significant connection between populations of “Amazonia Core”, which share high numbers of large blocks over a long distance (Fig. 4B). This sharing involves speakers of Cocama (a Tupí language) in Colombia, who share long IBD blocks with individuals from Wayku and in particular with individuals from the “LoretoMix” group in Peruvian Amazonia. The LoretoMix includes three Cocama speakers, and only these three individuals share IBD blocks with the Cocama from Colombia (marked with a red asterisk in Fig. 2A), despite a distance of more than 500km separating the two sampling locations. The strongest signal of relatedness is found between the neighboring Inga and Kamentsa populations from Colombia, who share numerous, long IBD blocks.

**Fig. 4.**
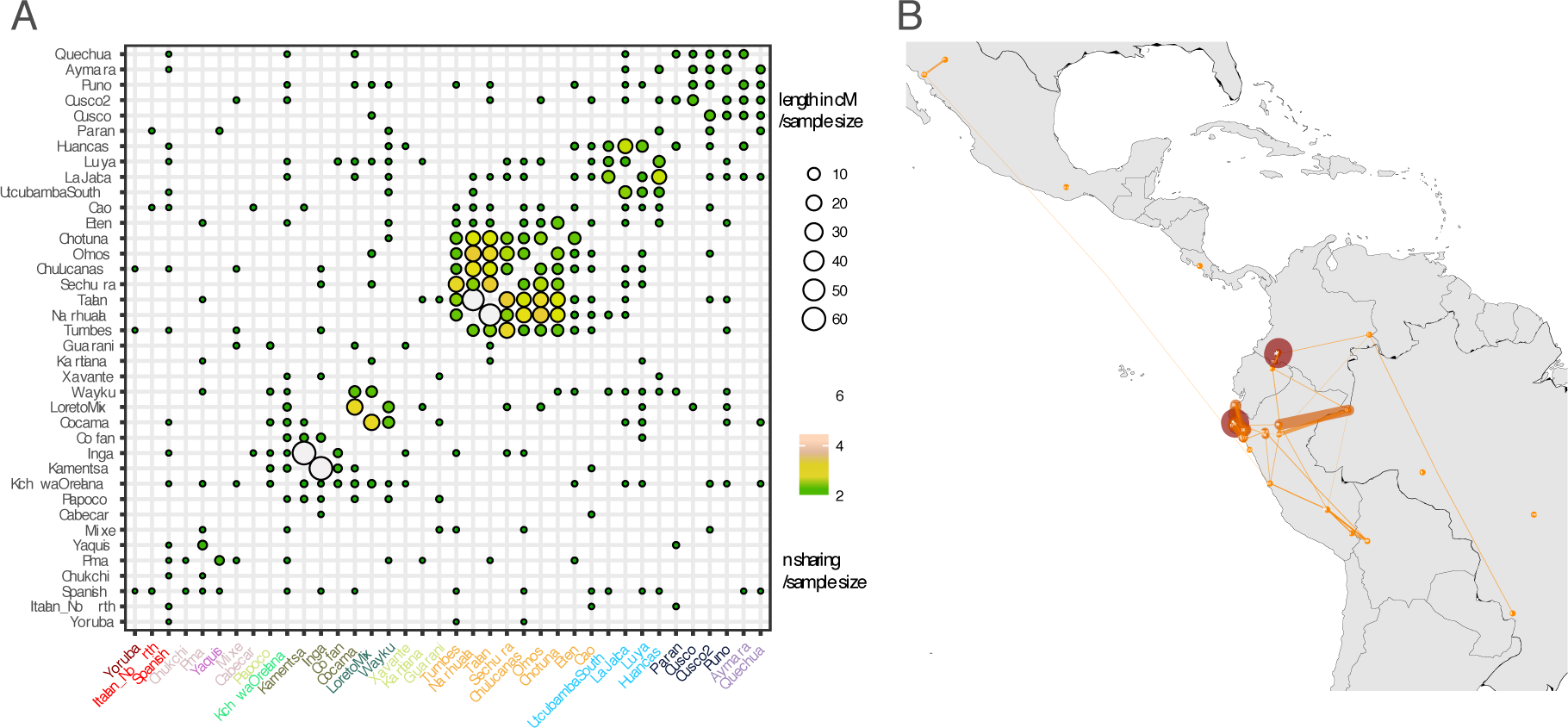
Results of the IBD sharing analysis. A. Symmetrical matrix of pairwise IBD blocks sharing, showing the total length and the number of occurrences adjusted by population size. Populations are ordered by ecoregion and color-coded as in Fig. 2. B. Map visualizing the connections between populations that share blocks with each other: thin yellow lines indicate the lowest levels of exchange, thick red lines the highest (adjusted for population size).

In North America, Yaquis share many long blocks with Pima (both speaking a language from the Uto-Atzecan family), at a distance of 250 km. Finally, numerous shorter fragments are found to be shared between Amazonia and the Andes, in particular between speakers of languages within the Quechua family: Kichwa Orellana and Wayku are connected with populations of the North-East and Central-Southern Andes.

A coancestry matrix generated with Fine Structure analysis (Fig. S7) clarifies the relationships at the individual level, by visualizing the intensity of blocks of shared ancestry, and confirms the results of the IBD analysis. In the central clusters, we find a specific connection between individuals from the North-East Andes and individuals from Cusco and Parán (Lima), which are also in proximity to another cluster that includes other North-East Andes individuals, together with Coast individuals from Cao. A separate cluster includes all of the remaining individuals from the Coast, and connects them with Southern Andean individuals (our sample from Puno and the Quechua and Aymara samples from the literature).

### Post-Columbian admixture with Europe and Africa

We examined the uniparental data (in terms of haplogroup frequencies) for a first overview of the proportion of Native vs. non-Native American ancestry in each population (Table S1). The typical Native American haplogroups for mtDNA are A2, B2, C1 and D1 (plus the less frequent D4h3 and X2a, the latter not present in our dataset), while for the Y chromosome they are Q and C3 ^1^. Fig. S8 shows that in most groups the frequency of Native American mtDNA haplogroups is 100%; the exceptions are groups from the Coast (Cao and Tumbes), which have a few individuals assigned to the African haplogroups L3 and L2 (Table S1, marked as “others” in Fig. S8). The frequency of Y chromosome Native American haplogroups is overall lower than the mitochondrial one, but it reaches 100% in all individuals in Amazonia Core, in the Central-Southern Andes and in Tallán, Narihuala, Chotuna and Eten in the Coast. Non-Native American haplogroups (mostly R, of European origin, but also E, potentially of African origin) predominate only in Chulucanas, Tumbes, Cao, and La Jalca (Table S1, marked as “others” in Fig. S8).

A supervised ADMIXTURE analysis was then performed to investigate the proportion of Native American ancestry per individual (Fig. S9). This analysis shows that several populations from all three ecogeographic domains display negligible proportions of European or African ancestry, confirming the results from the uniparental data. The proportion of Native American ancestry in the autosomal data, averaged per population, is roughly proportional to the average of female and male Native ancestry, with the latter being lower than expected in the admixed populations of the coast in particular (Fig. S10). The proportion of European ancestry is uniformly distributed among individuals only in North-East Andes. In all other populations that show evidence of European and African ancestry, the proportion of those ancestries varies widely at the individual level: this clearly suggests additional and more recent episodes of gene flow into these groups. For subsequent analysis of admixture (which are more robust for large sample sizes), populations were grouped according to similar Native American ancestry profiles, and outlier individuals were excluded (i.e. a single individual showing exceptional non-Native American ancestry among unadmixed individuals of the same population, as was the case in Sechura, Kamentsa and Cofán, and as indicated in the third column of Table S1). Furthermore, because the Colombian Inga clearly show structure with respect to their ancestry (Fig. 1), we separated the highly admixed individuals into an Inga_Admixed population, and merged the less admixed Inga individuals with the neighboring Kamentsa.

We used an f_3_ admixture statistic of the type f_3_ (Target; Source1, Source2) to confirm admixture events between Native American populations and European and African sources, where the target population is a South American population for which the ADMIXTURE results suggest European or African ancestry components. Source1 is a non-admixed Native American population (Xavante, Sechura_Tallan or Puno) and Source2 is either a European (i.e. Spanish) or an African population (i.e. Yoruba). Negative values of f_3_ confirm the signal of European admixture for a few populations of the coast, for the Mexican Yaquis, for all Andes North-East, for Cusco2, and for the Inga_Admixed (Fig. S11). African admixture appears in a subset of these populations, with the strongest signal in the Coast and in Inga_Admixed.

To further investigate the signal of recent admixture suggested by ADMIXTURE, we analyzed IBD blocks shared with Yoruba and Spanish sources. Sharing is detected in all three ecogeographic domains (Fig. S12). The largest number of blocks from the European source are found in Cao, Chulucanas, Tumbes, Cusco2, Yaquis, Inga Admixed, Luya and Utcubamba South. The pattern from the IBD sharing agrees with the profile from the supervised Admixture, with some exceptions: in Kichwa Orellana and Wayku, the IBD blocks imply more European ancestry than the ADMIXTURE results do.

To explore the intensity and timing of post-European contact in our selection of populations we employed two methods, which date admixture based on different aspects of the data: MALDER ^46^ and wavelet transform analysis (WT) ^47^. Both methods are applicable to admixture events involving more than two source populations. We again used Yoruba and Spanish as proxies for the African and European source populations, respectively. The results are summarized in Fig. 5.

**Fig. 5.**
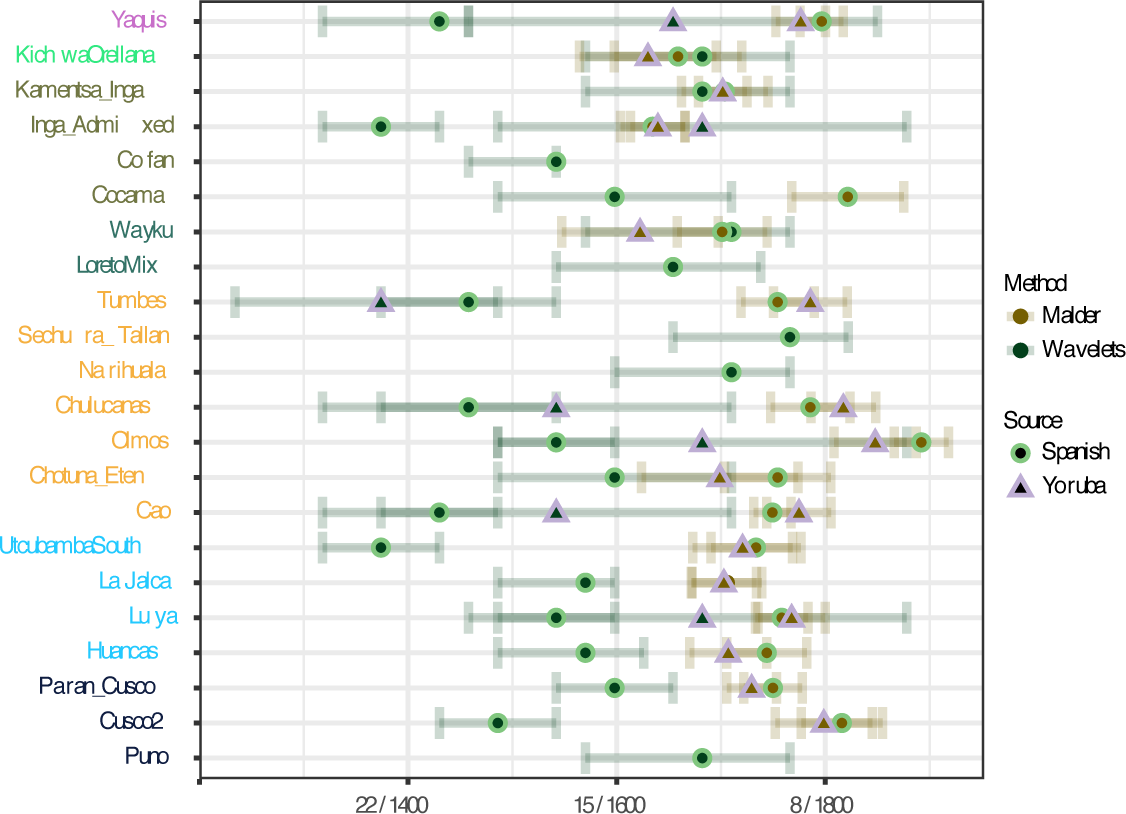
Admixture dates between European and African sources. Estimates of admixture are calculated with the MALDER and WAVELETS methods. Dates are expressed in generations ago and converted to calendar years using a generation time of 28 years.

Local ancestry along individual chromosomes was inferred using the RFMix method ^48^. With MALDER we ran the analysis to infer dates for both European and African admixture for all populations, regardless of the admixture proportions. With the WT-based method, meanwhile, for African admixture we inferred dates only if the proportion of African ancestry in a given population was over 1% (estimated based on RFMix). Overall, for most populations the dates inferred by WT are about 8 to 18 generations earlier than those inferred by MALDER. It has been shown previously ^49^ that this discrepancy between the two methods is expected in situations with continuous admixture or multiple pulses of gene flow from the same source, as MALDER is more sensitive to recent admixture events.

The dates inferred for European admixture are in most cases more recent than those for African admixture, reflecting an admixture history protracted through time for the European ancestry source. The most recent dates (for both African and European admixture) correspond to 7-8 generations ago for MALDER and 8-10 generations ago for the WT method. For MALDER, the most recent dates are found in the Yaquis, Chulucanas, Olmos and Cusco2. The older dates are found in Amazonia, in particular in our “Amazonia North” (Kichwa Orellana and Inga) and in Wayku, where the admixture is estimated to have happened between 1650 and 1700. Here the dates from MALDER and WT mostly overlap.

## DISCUSSION

We generated genome wide data with the Affymetrix Human Origins array for 176 individuals from 25 populations of North America and western South America, and analyzed these data together with published data from representative populations of the continent. Our strategy in collecting and analyzing the data can be summarized by three major objectives. 1) To investigate patterns of genetic diversity within and between the three main ecogeographical domains of western South America (the Andes, the Amazonia and the Coast), especially in understudied regions and in transitional environments. 2) To retrace past connections between and within the domains, and to evaluate to what extent the genetic landscape of South America was impacted by the last and largest complex societies of the pre-Columbian period, and by the distributions of indigenous language families. 3) To reconstruct the timing of admixture events from European and African sources in the centuries immediately *after* Columbus, and to identify differences in the chronology of such admixture in different populations within each of the three main domains.

For the first objective, we investigated Native American ancestry across the continent. One major Native American ancestry component is shared by all populations, as seen in the ADMIXTURE plot (Fig. S2, K=3, associated with the lowest Cross Validation error), in line with results from other living populations and from ancient DNA, which support a single major migration ^4^. This Native American component exhibits a marked population structure; the main signal of structure separates Amazonia from a shared Andes/Coast ancestry. Populations from North America were shown to have a distinct ancestry component in the ADMIXTURE analysis (Fig. S2), with a smaller differentiation between populations from southern North America (Yaquis and Pima) and Meso-/Central America (Mixe and Cabecar, Fig. S4). The diversification of these ancestry blocks from the initial single Native American gene pool does not bear traces of a north-south gradient of differentiation, or of serial founder effects, as the genetic distances between populations displays a radial structure rather than a sequential one (Fig. S4). A previously observed early diverging component similar to the Mixe ^10^ or the ancient Californian Channel Islands ^8^ is not captured by our data, which focuses more on South American genetic diversity. Amazonian ancestry is further split into two components: one more widespread in the Amazon Basin (here called “Amazonian Core”), the other corresponding to the piedmont populations of Ecuador and south-western Colombia (“Amazonian North”, Fig. 1 and 2). This latter component is strongly differentiated (it is one of the first ancestry components to appear in the ADMIXTURE analysis, at K=4 - Fig. S2), suggesting that it either diverged early in the migration process, or reflects stronger drift than in the rest of Amazonia. It is unclear if this ancestry could correspond to one branch of the earliest continental population structure, which were hypothesized from aDNA admixture graph reconstructions ^10^. This “Amazonian North” ancestry represents a previously unreported source of ancestry from a transitional environment: this region is in fact geographically close to the Northern Andes between Colombia and Ecuador, but its populations are traditionally associated with the Amazonian cultural domain. An early human settlement of Ecuador and northern Peru (between 16.0 and 14.6 kya) has previously been inferred from high resolution mtDNA data ^50^, in line with the archaeological record ^51^. Meanwhile, the presence of pockets of diversity in Ecuador and Colombia is paralleled by the presence of distinctive Native American lineages, such as Y chromosome haplogroup C3, otherwise rare in the continent ^52^. This haplogroup is also reported for other populations in Colombia ^53^. Interestingly, haplogroup C is found in the sample from Ecuador (Fig. S8, Table S1), too, but further analysis is needed to precisely identify the Y chromosome subhaplogroup and confirm the C3 affiliation.

Finally, populations from the Coast and the Central Andes (both north and south) show close genetic proximity to each other, as visualized by the PCA in Fig.s 2A and 2B and by the same ancestry component profile up until K=6 in Fig. S2). This strongly suggests a common origin and/or extensive contact, which may be associated with a coastal migration route and a colonization process from the coast inland into the highlands ^5,11,54–56^. Previous analyses have already noted the common history of these two regions, with first settlement from around 12,000 years ago ^11^.

For our second objective, on connections within and between domains, we explored signatures of demographic history and haplotype sharing patterns. The ROH variation profile of most populations from the Coast and the Andes (both North-East and Central-Southern) is consistent with a history of a relatively large population size, with some exceptions (Sechura, Narihuala, Cusco) that may have experienced isolation and drift only very recently (Fig. 3, Fig. S5). The long-term presence of large-scale state societies in the Andes and on the Coast can be expected to have promoted gene flow across wider geographical scales and merged previously structured populations, contributing to the higher genetic diversity of the current inhabitants. On the north coast of Peru, the Moche culture was one of the largest entities from the 1^st^ century CE, followed by the Chimú from the 12^th^ century ^57^. Their political influence over the coast would have promoted human connections that overcame the long stretches of desert that separate the main river valleys and the Humboldt current and wind regime that make long-distance seaborne trade difficult. In the Chachapoyas region (North-East Andes), a number of structured societies flourished from the 12^th^ to the 15^th^ century ^58^. In the Central-Southern Andes, the Wari and Tiahuanaco ‘Middle Horizon’ (c. 500-1000 CE) and especially the Inca ‘Late Horizon’ (c. 1470-1532 CE) established vast networks that mobilized and moved large labor forces for agricultural production (terracing, irrigation, raised fields), operated resource exchanges through camelid caravans, and resettled populations as explicit state policy ^57,59,60^. The impact of the Wari and Inca Empires is widely associated also with the diffusion of the two main surviving Andean language families, Quechua and Aymara ^61–63^.

The Coast and our two Andean sub-regions share a similar ancestry (as discussed above) and a similar history of large population size, but they are differentiated at a finer scale, with localized patterns of IBD segment sharing. Longer IBD blocks are shared almost exclusively within each of the three geographic macro-regions: Coast, North-East Andes and Central-Southern Andes, with the latter sharing fewer and shorter segments, suggesting more ancient contact over a large region, not sustained until recent times (Fig. 4). By contrast, the Amazonian populations in most cases have longer ROH blocks and overall high levels of consanguinity. This could reflect the model first proposed by ^14^: larger groups in the Andes vs. small, isolated groups in Amazonia. Nevertheless, by including more populations from a wider range of cultural and geographical backgrounds, we find exceptions to this model with some Amazonian populations characterized by a smaller number of larger blocks, belonging to group 1c and group 2 in Fig. S5. The populations of Amazonia therefore display different demographic histories rather than a uniform history of small sample size (according to the ROH profiles) and are connected by sharing of IBD blocks within the region. Moreover, Amazonian populations also show long-distance sharing of large and short fragments with the Andes and the Coast (Fig. 4), which is not consistent with the traditional portrait of isolation between Amazonian populations. This genetic diversity complements the evidence from other disciplines that the region was also home to dynamic, non-isolated population groups ^12^. In particular, the linguistic diversity of Amazonia includes not just language isolates but major, expansive language families, with far-reaching geographic distributions ^36^, reflecting long-range migrations of at least some speakers, and possibly major demographic expansions. There is also clear linguistic evidence for intensive interactions in convergence zones, and (more weakly) across Amazonia as a whole ^64^. We explored these potential connections by checking for gene-flow among speakers of the same language or language family. An interesting case is represented by the speakers of Cocama, a language of the Tupí family. The ROH profile for the Cocama of Colombia is lacking in long ROH segments (Fig. 3, Fig. S5), suggesting no recent bottlenecks or isolation. The analysis of shared IBD segments, which indicate shared ancestry through recent contact, reveals a long-distance connection between this population and geographically-distant populations of Peruvian Amazonia (Fig. 4). In particular, three Cocama speakers included in the LoretoMix sample from Peru are slightly different from the rest of the LoretoMix sample, and genetically closer to the Cocama of Colombia (Fig. 2A). Archaeological and ethnohistorical evidence indicates that the ancestors of the Cocama and Omagua were widespread in pre-Columbian times, inhabiting large stretches of the Amazon Basin and several of its upper tributaries ^65,66^. Thus, the sharing of IBD segments as well as the lack of long ROH in the Cocama could be explained by large, widespread populations that were connected in pre-Columbian times. Alternatively, more recent migrations could have carried the Cocama language between Colombia and Peru. Both time–frames and both scenarios suggest a parallel between genetic and linguistic history, with language acting as a preferential vector of population mobility.

Weak evidence for long-distance linguistic connections is observed not only within Amazonia, but also between Amazonia and the Andes. This is the case for Quechua-speakers of lowland Ecuador (Kichwa Orellana) and lowland north-eastern Peru (Wayku), who share relatively short IBD fragments with Central-Southern Andes and North-East Andes respectively. Previous results based on Y chromosome haplotype sharing did find a similar pattern of connections between lowland Quechua-speakers in Ecuador and north-eastern Peru, but did not find such long-distance connections with the Central and Southern Andes ^31,32^. These different results can possibly be justified by sex-biased gene-flow (i.e. less male mobility), which should be further investigated with denser sampling and high-resolution mtDNA genome sequences. Overall, this new genomic evidence points towards a demographic connection behind the diffusion of Quechua varieties not only in the southern highlands, as previously attested ^31^, but also in the north, across ecogeographic domains. The genetic signature reconstructed can inform the historical reconstructions for the diffusion of this language family.

Finally, for the third focus we explored the traces of post-colonial history and the impact of European mediated gene-flow (from Europe and from Africa through the slave trade) in the different ecogeographic regions. In our newly reported samples we find a high proportion of Native American ancestry, with some populations showing no detectable post-Columbian admixture in all three ecogeographic domains (Fig. 1). In parallel we detect a high proportion of Native American mtDNA and Y chromosome haplogroups (Fig.s S8 and S10). These results are in agreement with previous studies on ancestry proportions among Peruvian populations ^11,24^. Moreover, a high Native American ancestry proportion is even observed for the Coast, even though the traditional fishing/trading economy ^67^ might have been expected to introduce gene flow from other Native and non-Native sources. Importantly, our sampling strategy was guided to avoid individuals who self-reported any grandparent or parent of European, African or Asian descent, thereby introducing a first filter for recent admixture. Nevertheless, this strong Native American ancestry reveals the potential of undersampled regions of the Americas for further exploring pre-Columbian genetic history.

We used two different methods to estimate the date of admixture with European and African sources (Fig. 5). While simulations show that in simple one-pulse admixture situations both MALDER and WT-based methods perform equally well for both recent and older admixture times, the dates inferred by both methods are not concurrent in more complex admixture scenarios, involving either multiple pulses or continuous gene flow ^49^. MALDER is more sensitive to the most recent admixture event experienced by a population, while the WT method is more sensitive to older admixture events, and tends to give intermediate dates when there are multiple admixture pulses ^49^. Here, the WT method consistently returned older dates than MALDER, suggesting multiple and/or continuous admixture, and potentially also a signal of deep shared ancestry between Native Americans and Eurasians, as evidenced by studies of ancient DNA ^68^. The oldest WT dates may reflect the initial episode of admixture experienced by some populations during the earliest colonization by the conquistadors, historically dating to the mid-16th century. Of these populations, the majority have much more recent MALDER dates of 7-8 generations ago (around the beginning of the 19^th^ century), i.e. the populations of the coast, the admixed samples in the highland from Cusco (Cusco2), and the Yaquis of Mexico. It is reasonable to assume that the contact with Europeans began earlier in these regions: the recent admixture dates may be describing either continuous admixture or a second, more recent pulse of admixture (not necessary from European immigrants, but also from local *mestizos*). This would be compatible with the admixture profile of Peru as reconstructed by a recent study, where the major pulse of European admixture occurred during the 19th century, after the impact of the war of independence in Peru ^11,39^. Nevertheless, not all populations fit this profile of a recent admixture pulse: in “Amazonia North” and in North-East Andes (where La Jalca is the most isolated location), MALDER recovers older admixture dates, between 15 and 11 generations ago, which often overlap with the WT dates (Fig. 5). The admixed Inga sample fits the profile of “Amazonia North”, with relatively ancient admixture dates (as reconstructed by both methods), but the retrieval of a few long IBD blocks shared with Spanish (Fig. S12) and the overall high admixture proportions per individual (Fig. S2) suggest possible further recent admixture. These potential pockets of isolation from further pulses of admixture, which lasted for three centuries, indicate different historical patterns of integration, or a less continuous and ubiquitous gene flow from individuals who carry European ancestry, and is characteristically found in sampling locations within Amazonia. There, the admixture dates around 1650-1700 are in agreement with historical records of early intrusions of Spaniards (including missionaries) into Peruvian and Ecuadorian lowland rainforests ^32^.

Finally, studies of ancient DNA have shown that as much as one third of the ancestry in modern Native Americans could be traced to western Eurasia ^68^. Similarly, modern-day Europeans were found to be a mixture of three ancestral populations, one of which was a population deeply related to Native Americans ^69^. These findings imply that European (or more accurately, Eurasian) ancestry found in modern-day Native Americans may not have been acquired exclusively through admixture during the time of European colonization, but instead may reflect a much deeper origin. It is therefore possible that the WT method is picking up this signal of shared ancestry, which predates European colonization, and hence infers dates for some populations that are too early to be consistent with the first appearance of the conquistadors in the Americas, only after 1492.

Admixture with African sources appears with relatively older dates and shorter fragments (Fig. S12), as it did not continue through time with the same intensity as the admixture with European sources (possibly through *mestizos*). It is also possible that the African component was incorporated only through European-mediated gene flow, as individuals in our samples who carry African ancestry always carry European ancestry as well (Fig. S9). These cases indicate some degree of isolation over the last two centuries from the admixture that occurred during the periods of Spanish colonial rule (1530s to 1820s) and of slavery (which largely overlapped), and replicate historical records for African slavery in Peru ^70^. The proportion of African individuals in the population was at its peak before 1800, but declined rapidly in proportional terms during the nineteenth century. In the colonial period and indeed thereafter, the African population was heavily concentrated on the coast, where it was exploited for plantation agriculture. The decline after 1800 was proportional rather than absolute, as the proportion of both the European and the indigenous populations rise in censuses in Peru, for example. Finally, the incorporation of the African genetic component was typically mediated by European males, while during the period of slavery marriage between people of African descent was hindered by the Spanish colonial regime.

In conclusion, by targeting key regions of Western South America and focusing on high resolution SNP array data we are able to reveal demographic histories, ancient structure and recent connections between different ecogeographic domains. These connections are particularly interesting for the Amazonia, traditionally portrayed by genetic models as a region of small isolated communities.

We also note how the widely analyzed population samples from the literature, e.g. the Karitiana and Xavante, exhibit high levels of genetic drift in comparison to our newly generated dataset — see the analyses of population relationship (Fig. S4) and of within population diversity (Fig. 3, Fig. S6). It is important to stress that inferences on Native American prehistory should not be drawn exclusively from such divergent populations with many closely-related individuals, but should instead include more diverse populations from different regions and different cultural and demographic backgrounds, in order to capture the diversity of the continent ^40,71^.

## METHODS

### Sample collection

Samples were collected during anthropological fieldwork expeditions by RB and CZ (Ecuador, 2007), LA (Colombia, 2012), CB, RF, JRS, and OA (Peru, 1998, 2007, 2009, 2014, 2015), and AAC and RSO (Mexico, 2016). The sampling collection and the project were approved by the Ethics Committee of the University of San Martín de Porres, Lima (Comité Institucional de Ética en Investigación de la Universidad de San Martín de Porres — Clínica Cada Mujer, Ofício No. 579-2015-CIEI-USMP-CCM, 12/05/2015), the ethics committee of the Universidad del Valle in Cali, Colombia (Acta No. 021-010), the Ethics Commission of the University of Leipzig Medical Faculty (232/16-ek), the Ethics Committee of the University of Jena (Ethik-Kommission des Universitätsklinikums Jena, Bearbeitungs-Nr. 4840-06/16), the Research Council for Science and Technology (Consejo Nacional de Ciencia y Tecnología - CONACyT, grant # 69856; Instituto Nacional de Ciencias Médicas y de la Nutrición Salvador Zubirán Ref.: 1518), and the National Commission for Scientific Research of the Mexican Institute for Social Security (IMSS; CNIC Salud 2013-01-201471). All methods were performed in accordance with the relevant guidelines and regulations, and in compliance with the rules of the Declaration of Helsinki. The samples analyzed in this study represent only a small fraction of the population living in the target regions of Mexico, Peru, Colombia and Ecuador, and so is only partially representative of the complex demographic history of these regions and of their inhabitants’ ancestors.

Details of the sampling collection and DNA processing are reported in ^12^ for the four Colombian population samples and in ^31^ for the Peruvian samples from Luya, La Jalca, Huancas, Utcubamba South (department of Amazonas) and Wayku (department of San Martín). The samples identified as “Cusco2” correspond to individuals who were sampled in the urban districts of San Sebastián and San Jerónimo (Cusco, Peru); details of the sampling are described in ^72^. Samples identified as Ecuador Kichwa were previously analyzed in search for a genetic variant associated with lipid metabolism ^73^. The other population samples have not been previously reported or described.

Samples from the population named “LoretoMix” include three speakers of Cocama (a language of the Tupí family), one of Chamicuro (Arawak), one of Shawi (Cahuapanan) and two of Muniche (a language isolate). These samples were collected in various locations within the department of Loreto in the Amazonian region of north-eastern Peru, and merged into one population after verifying their genetic affinity. The population samples from Cusco and Cusco2 consist of speakers of southern Quechua. The population sample labelled Puno (department) is made up of five speakers of southern Quechua and two of Aymara, collected on the islands of Lake Titicaca and merged into one population after verifying their genetic affinity. Parán is a community located in the highlands of the department of Lima, who speak Spanish. The population samples from the north coast of Peru include participants from rural areas and fishing communities who speak Spanish. The various population samples have been identified by the names of the towns or provinces where the samples were collected. Samples from the population named Kichwa Orellana include individuals sampled from the rural parish of San José de Guayusa, in the province of Orellana, in the Amazonian lowlands of Ecuador. The community speaks a variety of lowland Kichwa (the local name of Quechua), and includes individuals who recall relationships with Shuar communities from southern Ecuador. Samples from Yaquis were collected in the state of Sonora in north-western Mexico in the community of Tórim. People living there continue the culture and traditions of the Yaqui Nation and speak Yaqui, a language of the Uto-Aztecan family. The Mexican sample was included as a comparative source of genetic diversity from indigenous North America.

The samples have been subdivided into seven groups by country and macro-region (Mexico, Colombia Amazonia, Ecuador Amazonia, Peru Amazonia, Peru North-East Andes, Peru Central-Southern Andes, Peru North Coast). Individual information with details on the population, language spoken and geographic grouping is listed in Table S1. The sample locations for each population are shown in Fig. 1 and in more detail in Fig. S1.

### Data generation and screening

The DNA samples were screened and quantified with a Nanodrop spectrophotometer and Qubit fluorometer, and visually assessed by gel electrophoresis at the laboratory of the Department of Archaeogenetics of the Max Planck Institute for the Science of Human History in Jena. Sample genotyping was performed by ATLAS Biolab in Berlin on the Affymetrix Axiom Human Origins array ^41^. Genotyping data were processed using Affymetrix Genotyping Console v4.2.0.26. In total 188 samples were genotyped and genotyping call rates were >98.5% for all SNPs. The final dataset comprised 633994 SNPs. PLINK v1.90b5.2 ^74^ was used to calculate the inbreeding coefficient F and Pi_Hat values (degree of relatedness) between pairs of individuals, filtering for minimum allele frequencies of 0.05. One individual with a high F value was excluded and only one individual was kept in eight pairs with PI_Hat > 0.5. The same sample was included twice for cross-reference (CH008): 700 positions differ between the two, corresponding to an error rate of 0.1% in the genotyping. One duplicated sample was found, probably due to mislabeling. The final screened dataset consists of 176 individuals which were included in the analysis.

### Data availability

To access the genotyped data, researchers should send a signed letter to C.B. containing the following text: ‘‘(a) I will not distribute the data outside my collaboration; (b) I will not post the data publicly; (c) I will make no attempt to connect the genetic data to personal identifiers for the samples; (d) I will use the data only for studies of population history; (e) I will not use the data for any selection studies; (f) I will not use the data for medical or disease-related analyses; (g) I will not use the data for commercial purposes.’’

### Uniparental markers

Mitochondrial haplogroups were assigned with Haplogrep ^75^, limiting the call to major haplogroup nodes, given the uncertainty arising from the low number of mtDNA SNPs included in the Human Origins Array. Y chromosome haplogroup assignment was performed with the yHaplo software ^76^. Data was cross-checked with available published mtDNA and Y chromosome data for the same individuals, assigned via direct genotyping/sequencing in previous studies ^12,31,53^: the SNPs available allowed the correct macro haplogroup to be detected in 97% of cases.

### Merging

The newly generated dataset was merged with published Human Origins data from ^9,41,69^, selected to include populations representative of North and South America and of post-colonial African and European ancestry (Yoruba, Spanish and Italian North were chosen for these analyses). Not all samples or populations were used for all analyses, as described for each analysis. Merging was performed with the mergeit command in AdmixTools ^41^. A total of 597,569 SNPs were left after merging.

### Admixture analyses

We used the ADMIXTURE software ^77^ to infer individual ancestry components and admixture proportions, after performing LD pruning with PLINK. The LD pruning included the following settings, which define window size, step and the *r*^2^ threshold: –indep-pairwise 200 25 0.4 ^49^, leaving 232,755 SNPs.

We ran ADMIXTURE for values of K from 3 to 12, with 10 runs per K. We checked for consistency between runs and used the cross-validation procedure implemented in ADMIXTURE to find the best value of K. Population outliers such as Pima, Karitiana and Cabécar were excluded from this analysis — only Xavante was kept as a reference for Amazonian populations. Supervised Admixture (K=3) was performed to estimate the proportion of African, European and Native American ancestry per individual, keeping Yoruba, Spanish and Xavante (the latter known to be mostly unadmixed with European and African sources) as proxies for the parental groups.

We calculated f_3_ statistics as a formal test for admixture, using the same European/African parent populations as suggested by the results of the ADMIXTURE analysis, and with three unadmixed Native American populations with sample size larger than 6 (Xavante for the Amazonia, Puno for the Andes, Tallan together with Sechura for the coast). The qp3Pop command from the AdmixTools package was used to run f_3_. For each target population, the highest f_3_ value was kept (corresponding to the best choice of Native American parental population among the three proposed).

### Principal Component Analysis

Principal Component Analysis (PCA) was performed with smartpca of the Eigensoft package^78^. Different runs were performed with the whole dataset and with a subset of SNPs ascertained in the Karitiana (Panel 7 as identified by ^41^), consisting of 2,545 SNPs. SmartPCA was also used to calculate F_ST_ distances between populations, which were used to generate Neighbor-Joining trees in R with ape ^79^.

### Runs of Homozygosity and consanguinity

Runs of Homozygosity (ROH) blocks were identified with PLINK with default settings ^80^. We divided ROH in each individual into two categories, long ROH (>1.6 Mb) and short to intermediate ROH (<1.6 Mb), based on the classes defined by ^43^. We calculated the summed total length of ROH for each bin category for each individual. Long ROH were then further divided for a total of six bin categories and resulting ROH profiles were used to make demographic inferences following ^34^.

### Phasing and Identity by Descent analysis

BEAGLE v 5.0. ^81^ was used to phase the data. Before phasing, invariant sites were removed and the data was split into single chromosomes. Identity By Descent (IBD) blocks were inferred with refinedIBD ^82^. Three runs of phasing and IBD detection were performed for each chromosome. The runs were then merged and gaps were removed with the utility provided, allowing a maximum gap length of 0.6 cM and at most 1 genotype in an IBD gap that is inconsistent with IBD. Only blocks with a minimum length of 2cM and LOD score >3 were retained for the analysis, to avoid spurious calls and errors in block merging ^82^. The number of shared IBD blocks between pairs of populations was adjusted for sample size, by dividing by the product of the number of individuals in population 1 and population 2 in the pairwise comparison. Population pairwise sharing was considered only when more than one IBD block was retrieved, to further filter out spurious population connections. For the intra-continental comparisons, we considered fragments larger than 5cM, a threshold used in previous work that has found that shorter fragments are ubiquitously shared across the entire continent ^11^.

### Chromopainter and Finestructure

FineStructure v2 ^83^ was also used to identify ancestry blocks resulting from shared descent. Phased data were analyzed with Chromopainter to infer a co-ancestry matrix, followed by FineStructure for population clustering, following the standard process as described in the manual. A coancestry heatmap, a dendrogram and PCA plots were generated with the R commands provided in the package.

### Dating Admixture events

Dating of admixture events was performed via two approaches. For dating with MALDER^46^, populations with low sample sizes and similar levels of admixture (as estimated with the Supervised ADMIXTURE analysis) were combined, and outlier individuals with exceptionally high level of admixture were excluded from populations in which admixture was otherwise low or absent (Sechura, Cofán, Kamentsa - see Table S1). MALDER assesses the exponential decay of admixture-induced linkage disequilibrium (LD) in a target group, allowing for multiple admixture events (in this case for African, European and Native American sources). We ran MALDER with Yoruba, Spanish and three Native American parental populations, following the f_3_ analysis scheme. Only admixture cases supported by p value<0.05 and Z score>3 were considered. For each population and for each of the Native American parental groups that passed this filtering, the pair with the highest Z score was kept.

As a second approach we used RFMix ^48^ to estimate local European, African or Native American ancestry along individual chromosomes, and then applied wavelet-transform analysis to the output, and used the WT coefficients to infer time since admixture by comparing the results to simulations, as described previously ^47,84^.

Time in generations ago was converted to calendar years assuming a generation time of 28 years ^85^.

### Data visualization and source code

All data visualization was performed in R using packages developed by ^86–88^ and in-house scripts. The full detail of the command line setup and R scripts can be found at https://github.com/chiarabarbieri/SNPs_HumanOrigins_Recipes/

## Author contributions

C.B. designed research and analyzed the data; C.B, R.B., and A.M.T-A. performed laboratory analysis; C.B., R.B., L.A., J.R.S., O.A., C.Z., A.A-C., R.S-O. and R.F. collected samples; I.P. performed admixture analysis; R.G. and M.S. provided laboratory resources and reagents; P.H., K.K.S, M.S, I.P. and L.F-S supervised the research; C.B. wrote the paper with inputs from all the coauthors.

## Acknowledgments

We thank all the study participants and fieldwork assistants from Colombia, Ecuador, Mexico and Peru for making this study possible. The study was supported by a Wenner-Gren postdoctoral grant (Gr. 9395) to C.B., by the University Research Priority Program of Evolution in Action of the University of Zurich to C.B. and K.K.S., and by MEXT Kakenhi 18H05080 to K.K.S. L.F.S. was supported by a U.S. National Science Foundation grant (NSF: A15-0187-001). L.A. was supported by a graduate grant from COLCIENCIAS. We thank David Reich and collaborators for providing a formatted dataset of published data used for population comparisons and Adrian Pearce for historical contextualization of European and African admixture in Peru.

---

**Fig. S1.**
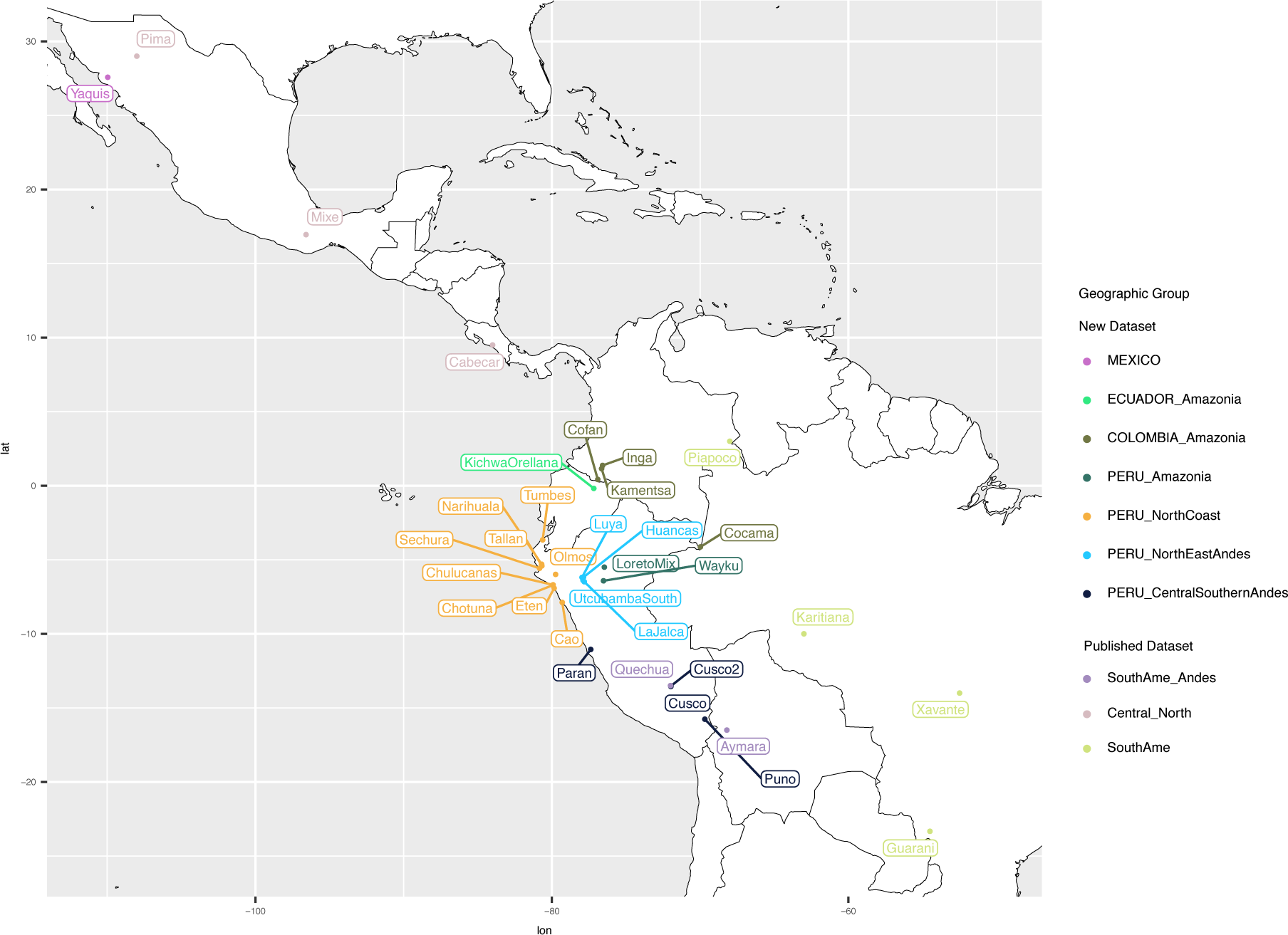
Approximate location of the population samples.

**Fig. S2.**
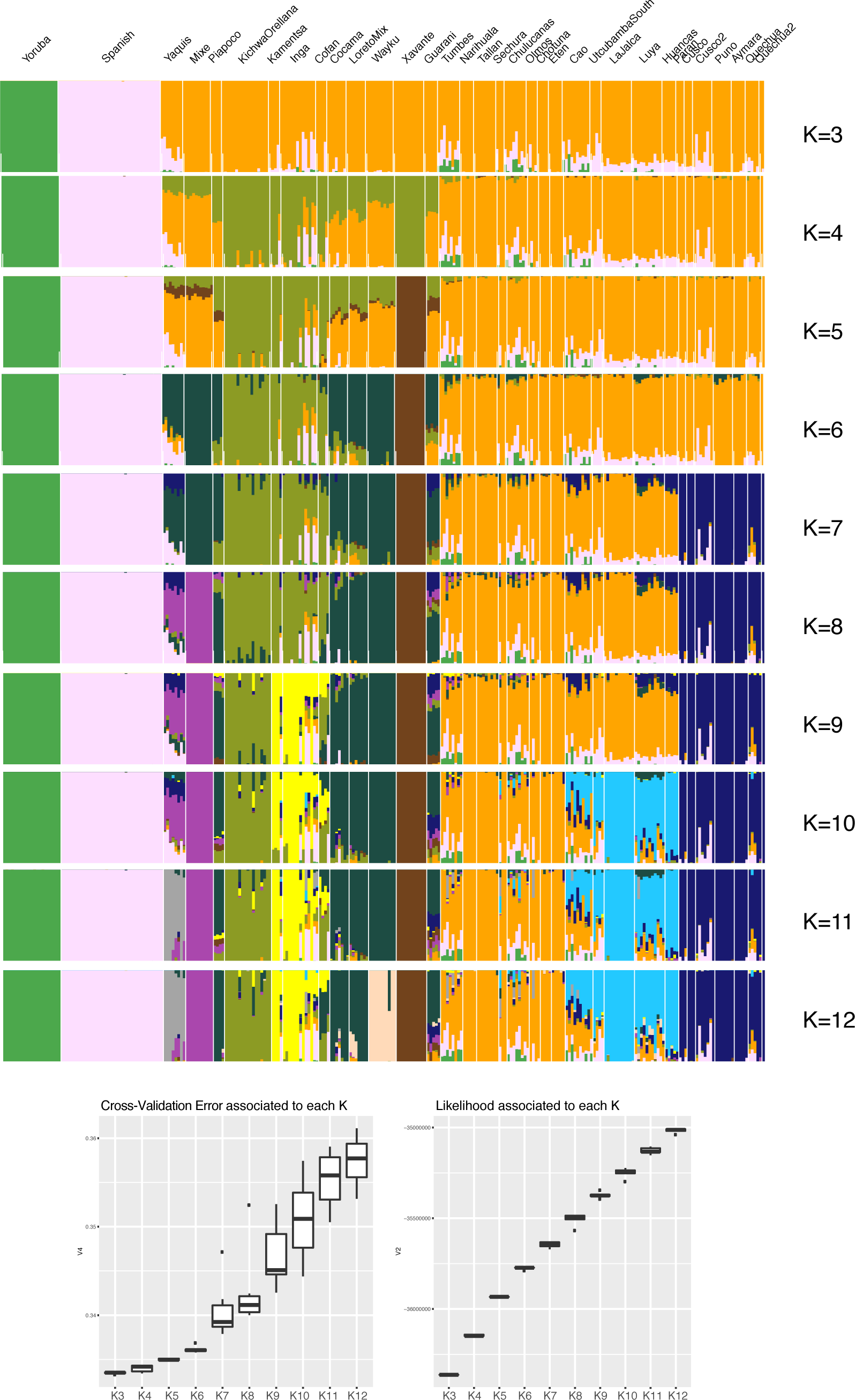
Admixture results for K between 3 and 12, with cross reference validation errors.

**Fig. S3.**
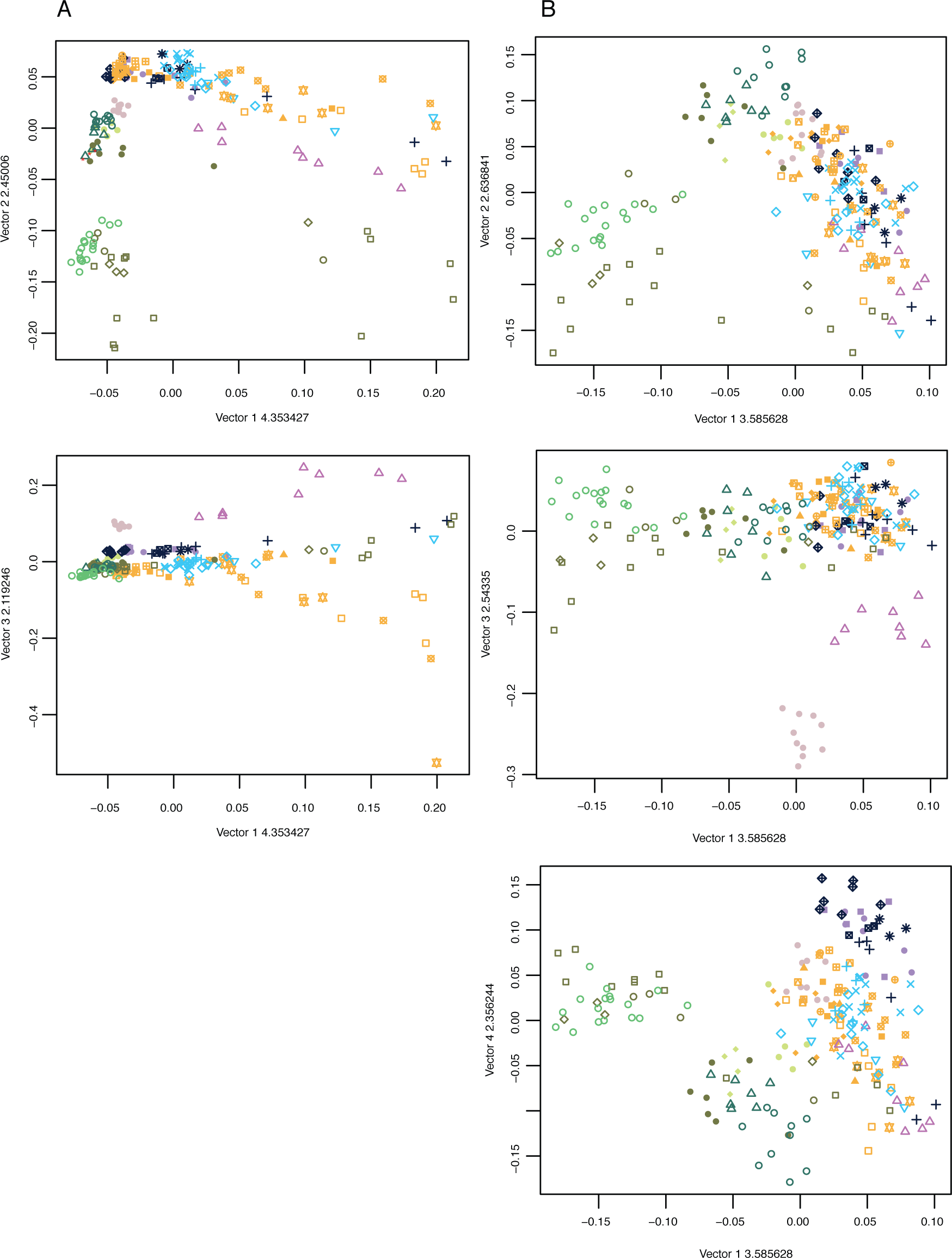
Principal Component Analysis of the newly reported samples together with representative populations from North and South America. A. whole SNP set, first three components. B: reduced SNP set ascertained for Karitiana (Panel 7, see Methods), first four components. Color legend corresponds to geographic grouping.

**Fig. S4.**
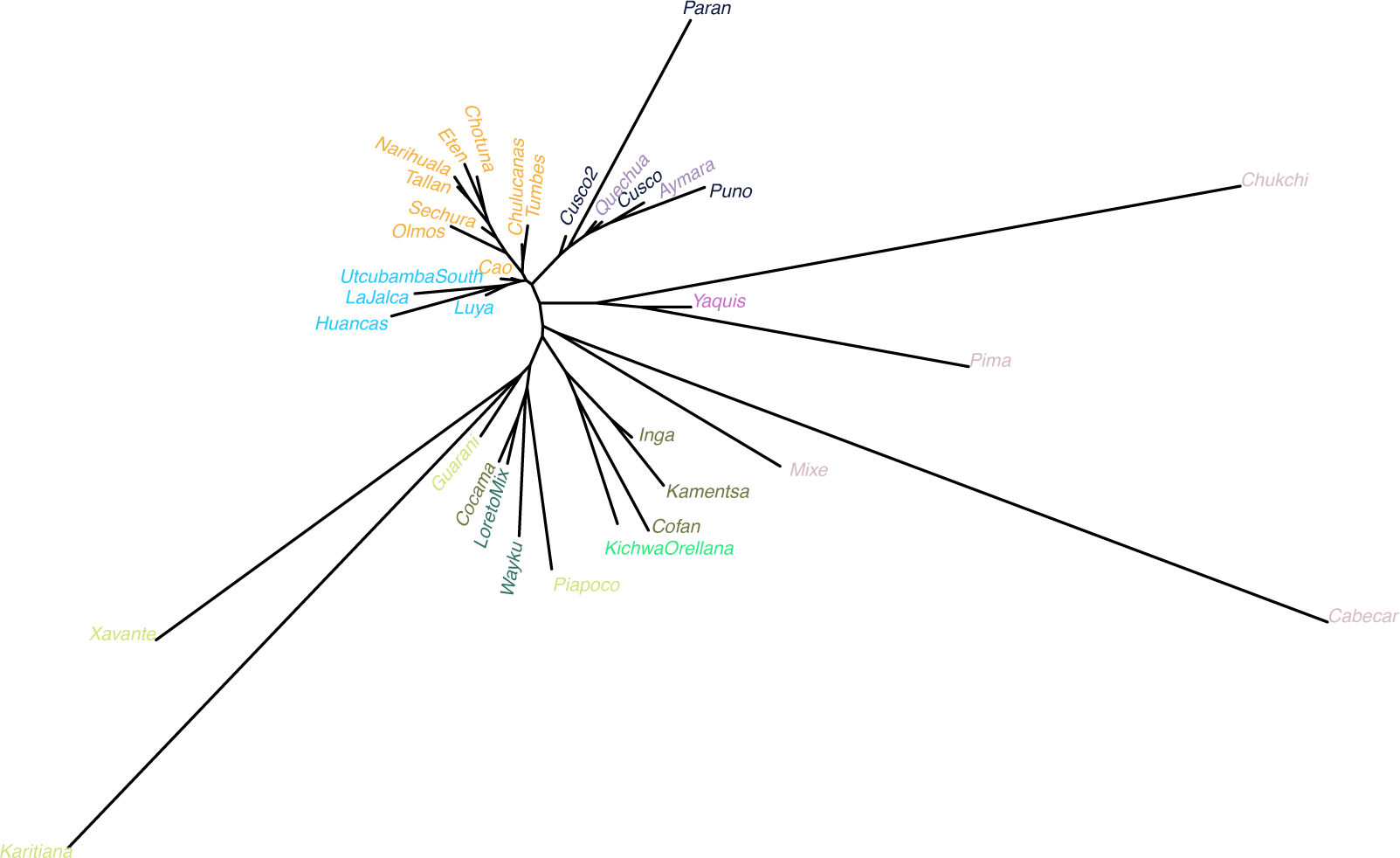
Neighbor Joining tree displaying F_ST_ distances between populations. Populations are color-coded as in Figure 2.

**Fig. S5.**
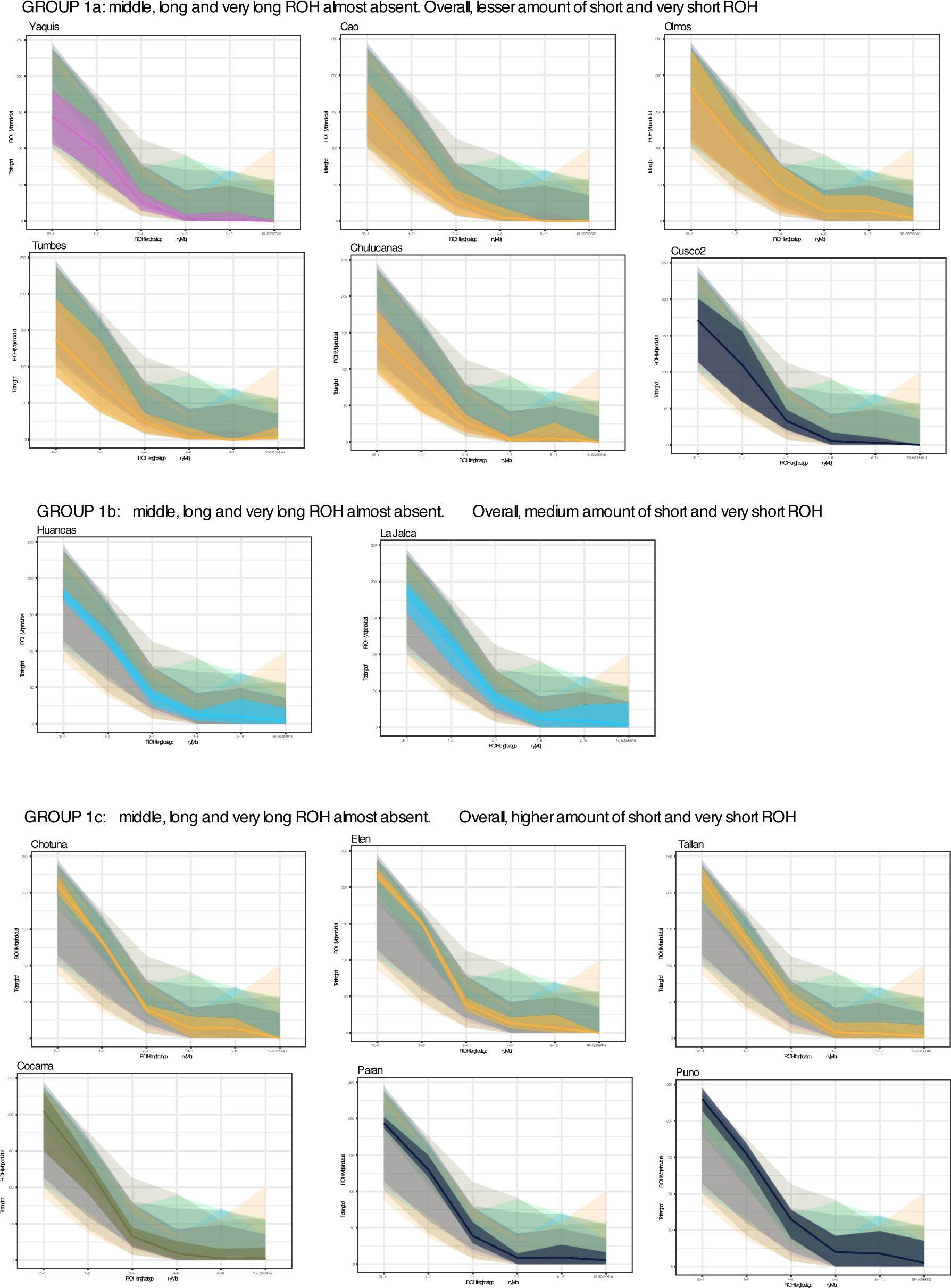
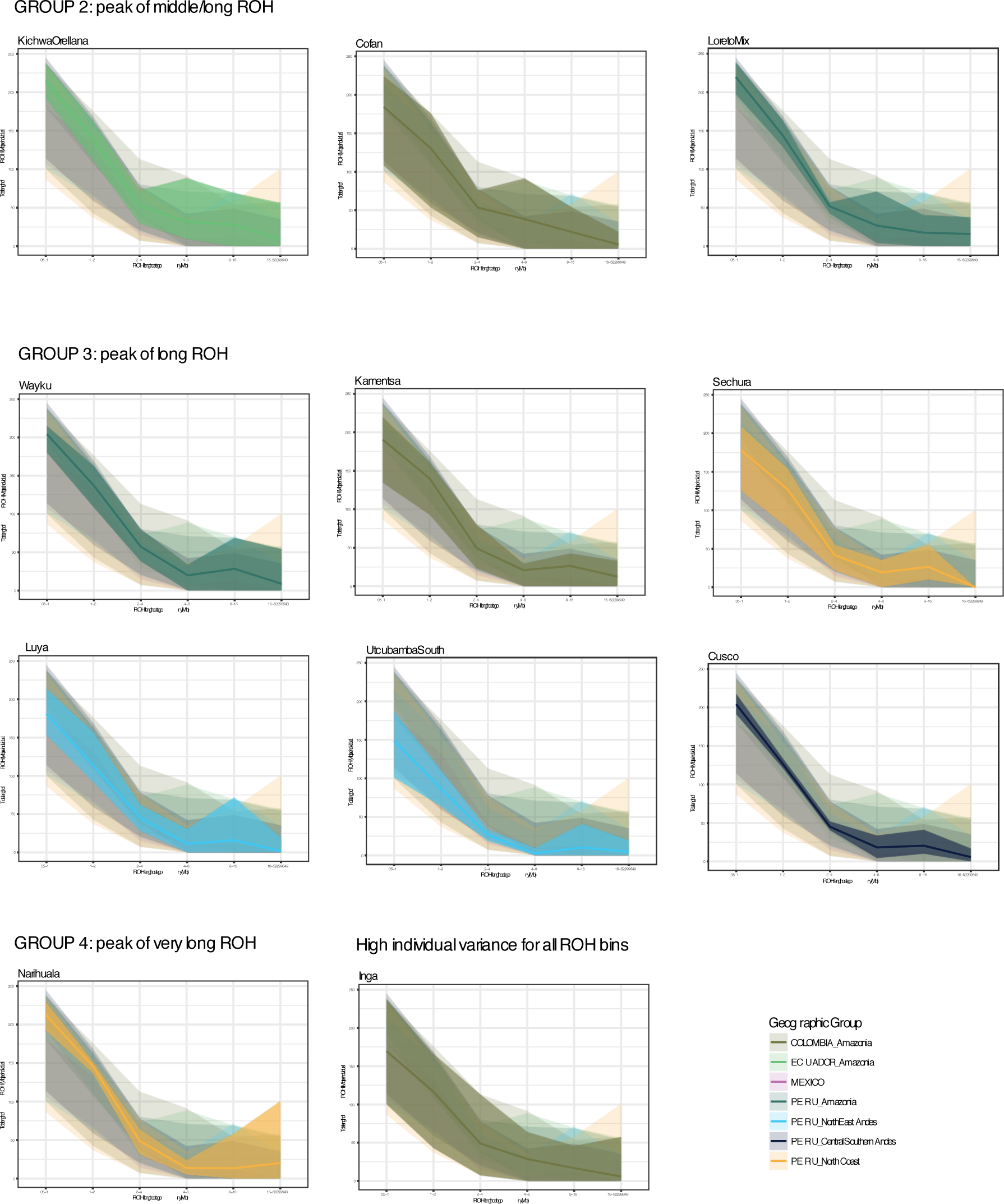
ROH length classes profile per population. Each plot shows the variance of total length of ROH per each individual, binned for the six classes proposed. Populations are grouped according to the similarity in their profile.

**Fig. S6.**
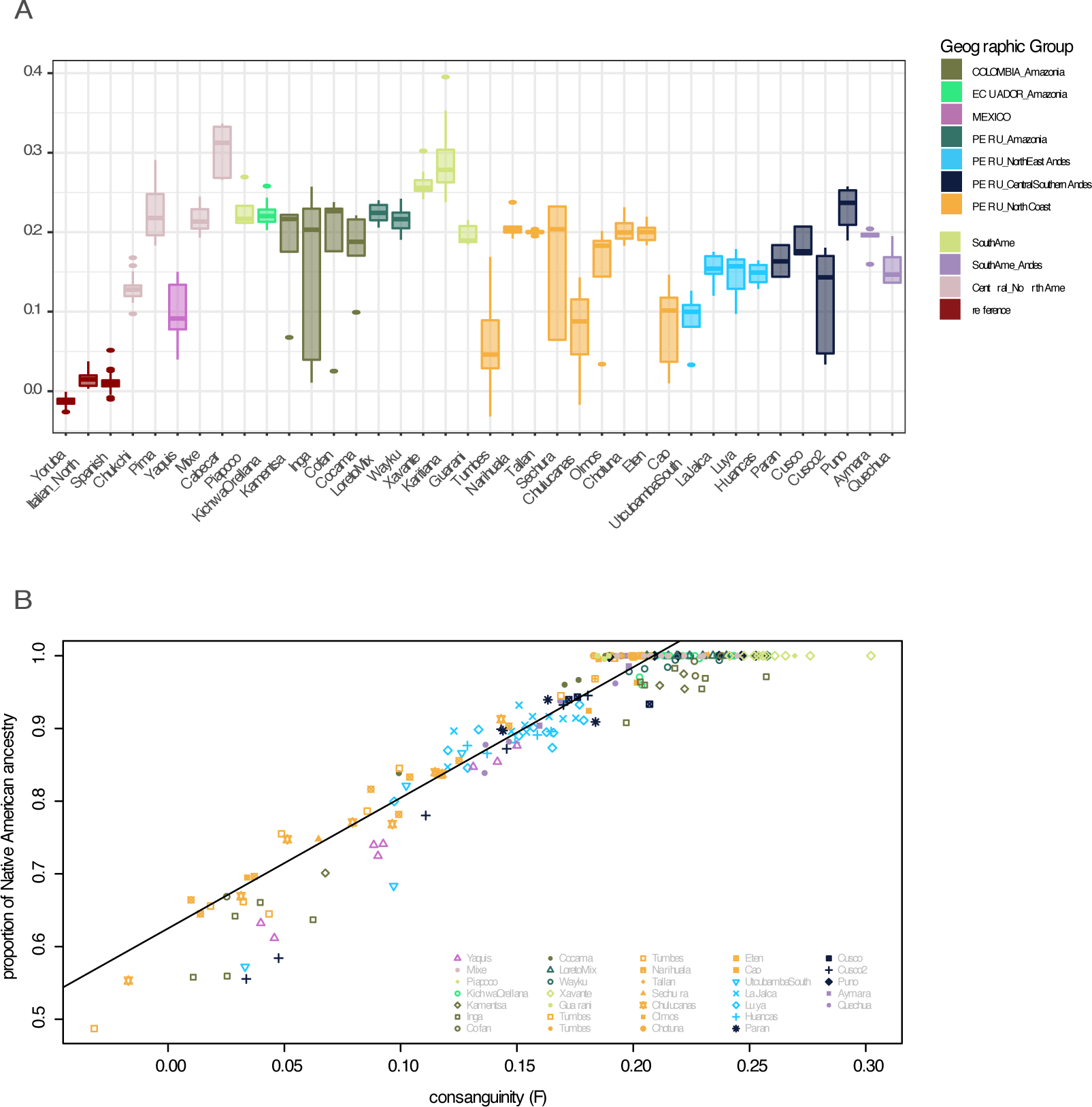
A. Distribution of measures of consanguinity (F) per individual for each population. B. correlation between consanguinity and percentage of Native American ancestry, calculated with Supervised Admixture Methods.

**Fig. S7.**
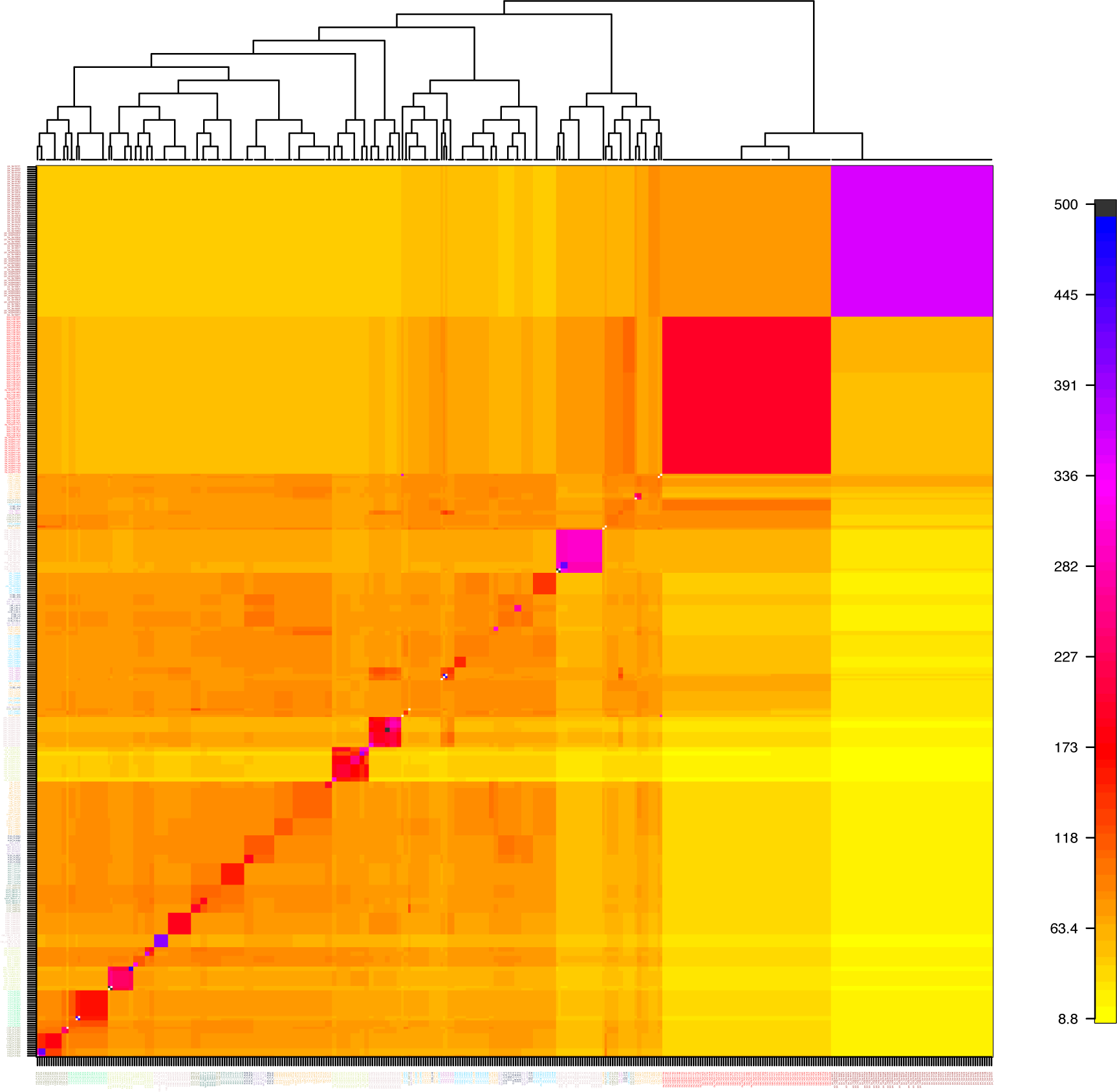
Population averaged coancestry matrix and individual dendogram generated with FineStructure.

**Fig. S8.**
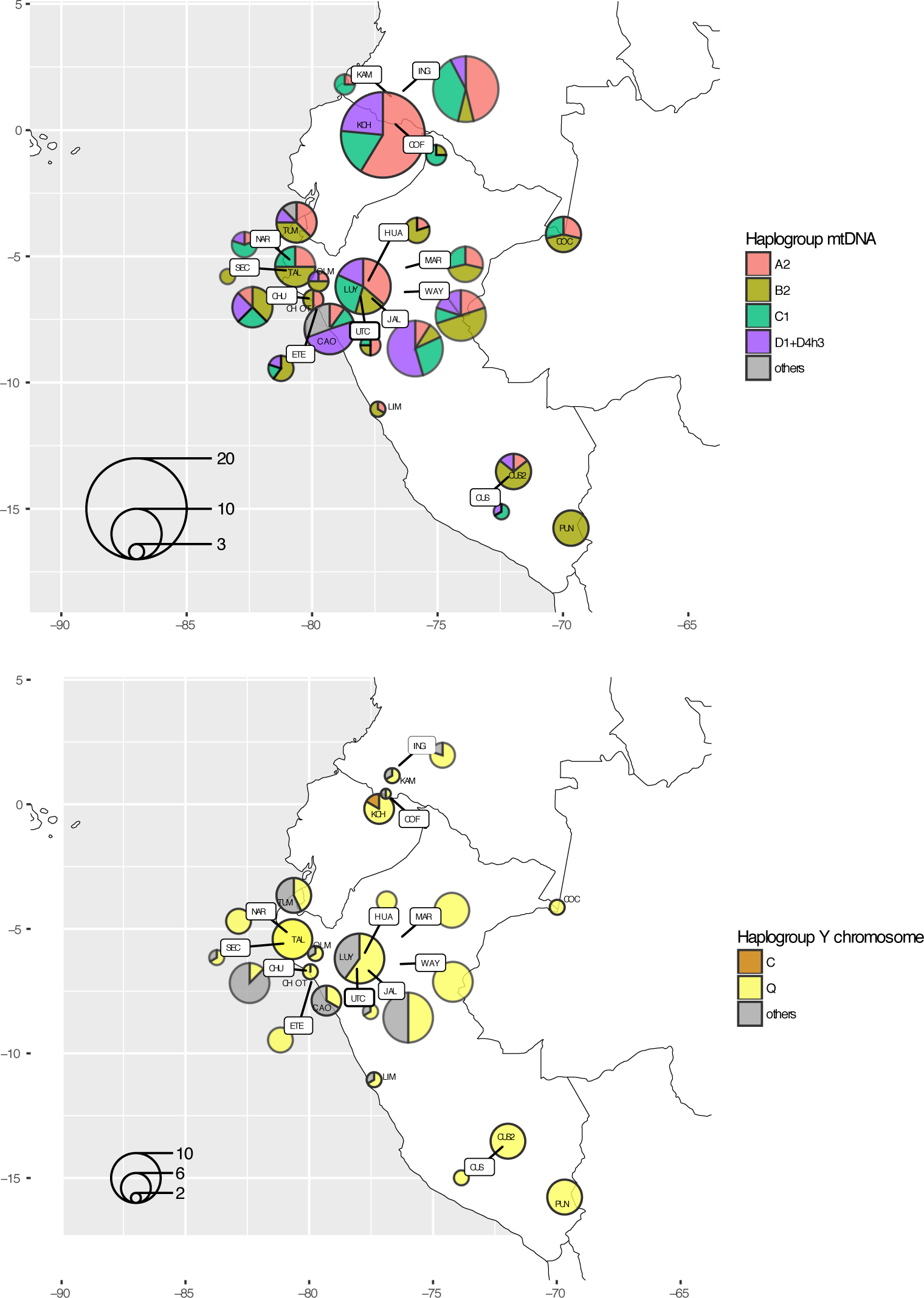
Pie Charts showing the haplogroup composition of each population sample, with approximate location on the map. A. mtDNA haplogroups; B: Y chromosome haplogroups. Note that the sample size is smaller for the Y chromosome haplogroup, because only the male individuals are considered.

**Fig. S9.**
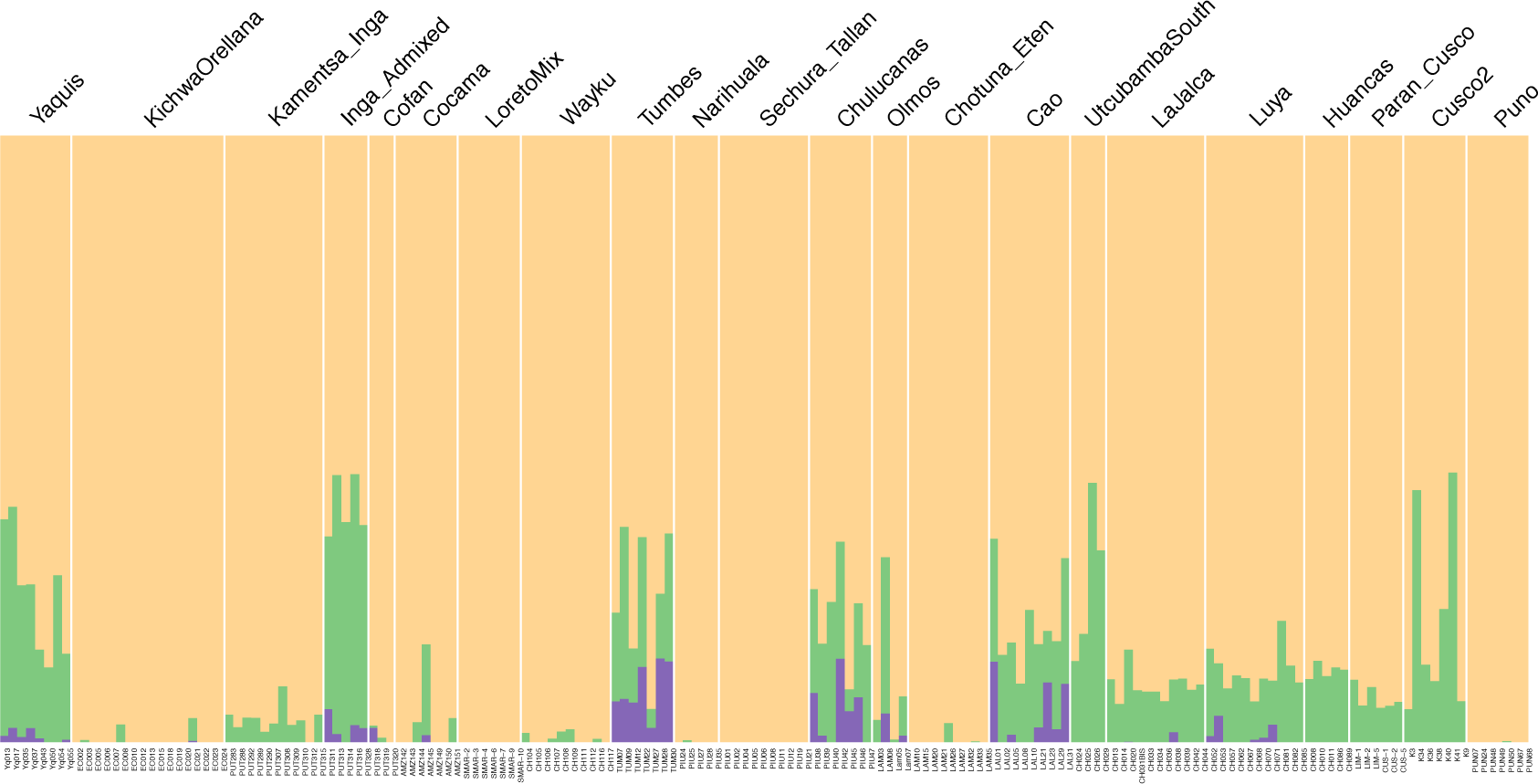
Results of supervised Admixture analysis at K=3.

**Fig. S10.**
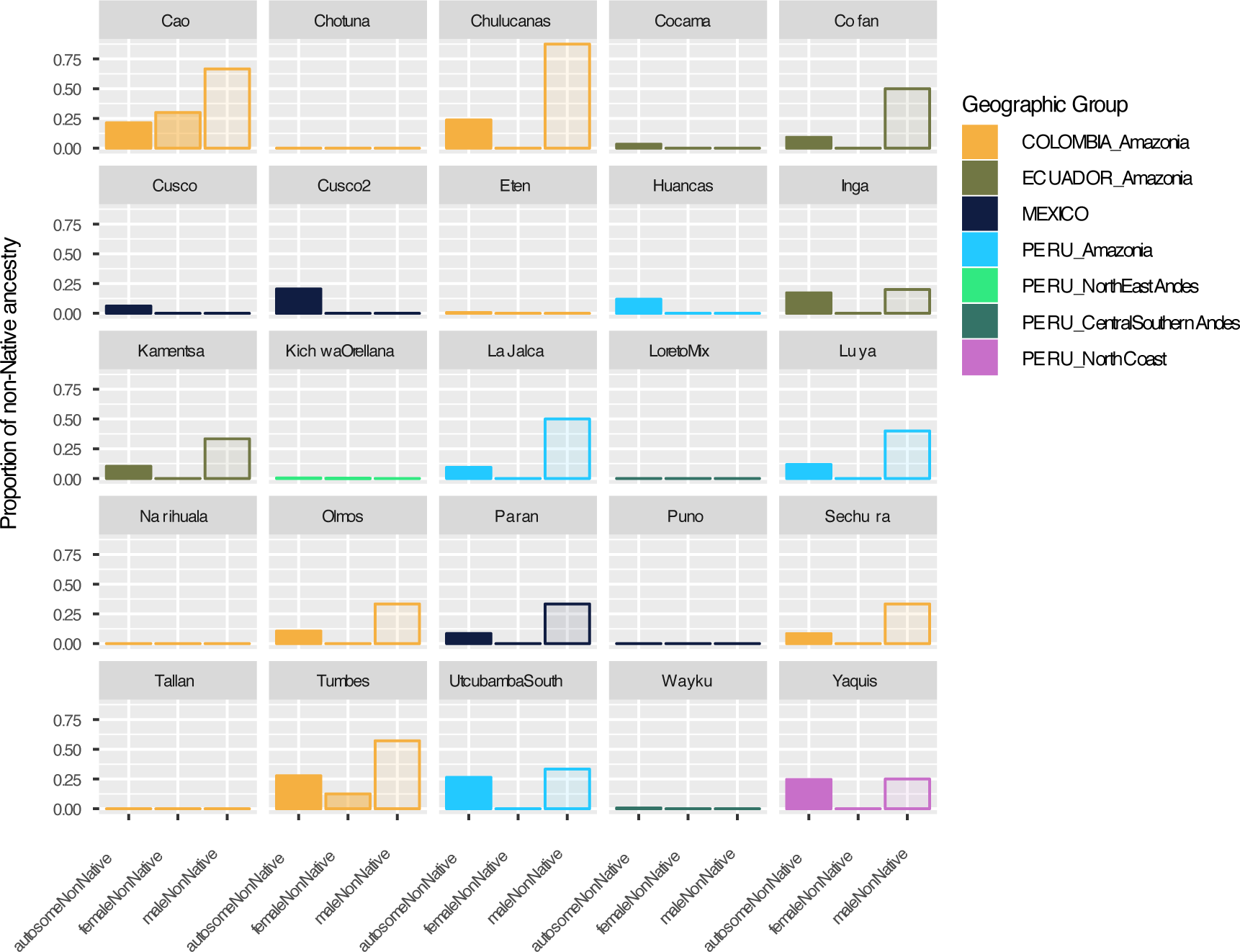
Proportion of Non-Native American ancestry per population, including autosomal (averaged over individuals, from Supervised ADMIXTURE analysis), Y chromosome and mtDNA (from haplogroup frequencies).

**Fig. S11.**
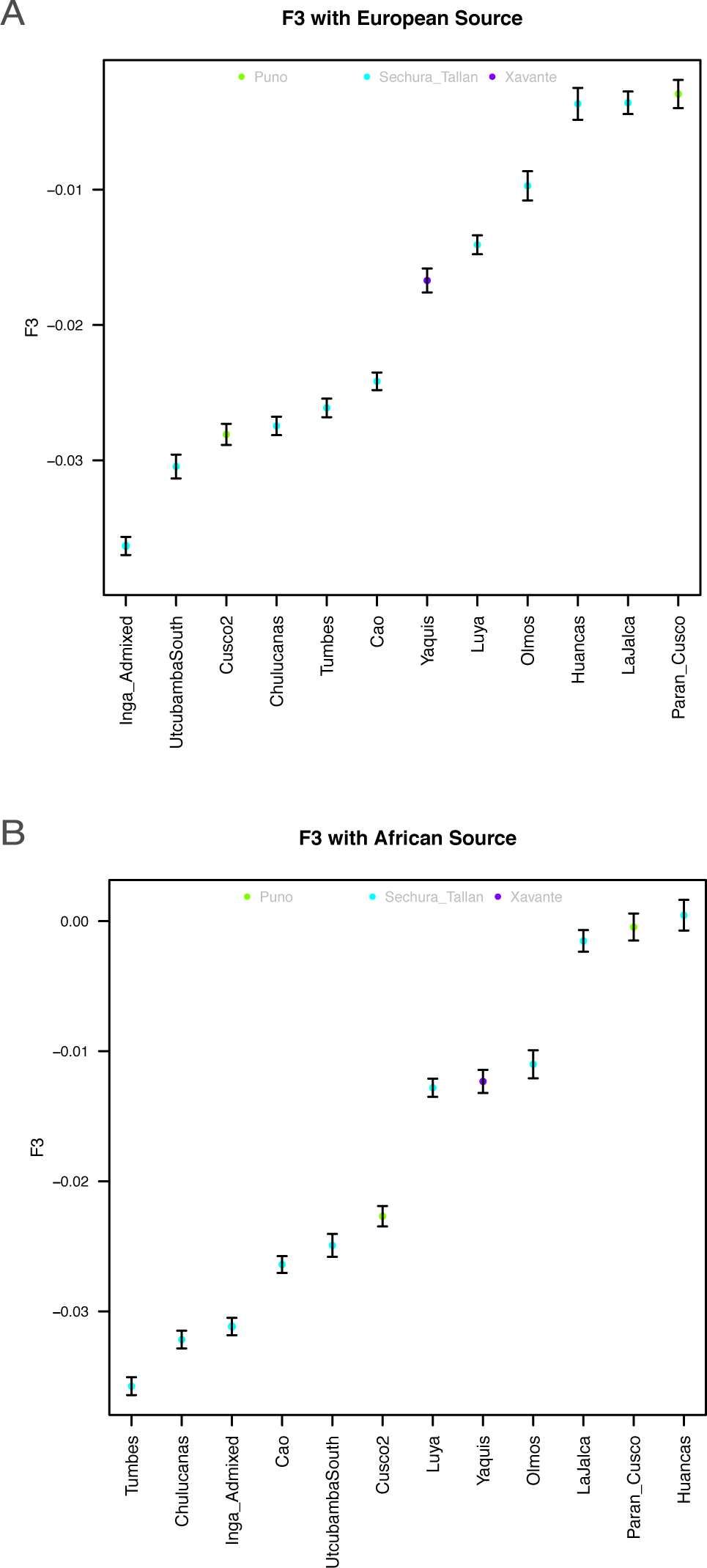
Results of the F3 analysis for A. European (Spanish) and B. African (Yoruba) admixture, ordered by the lowest (strongest admixture signal) to the highest. The color of the dot represents the source population which returned the lowest F3.

**Fig. S12.**
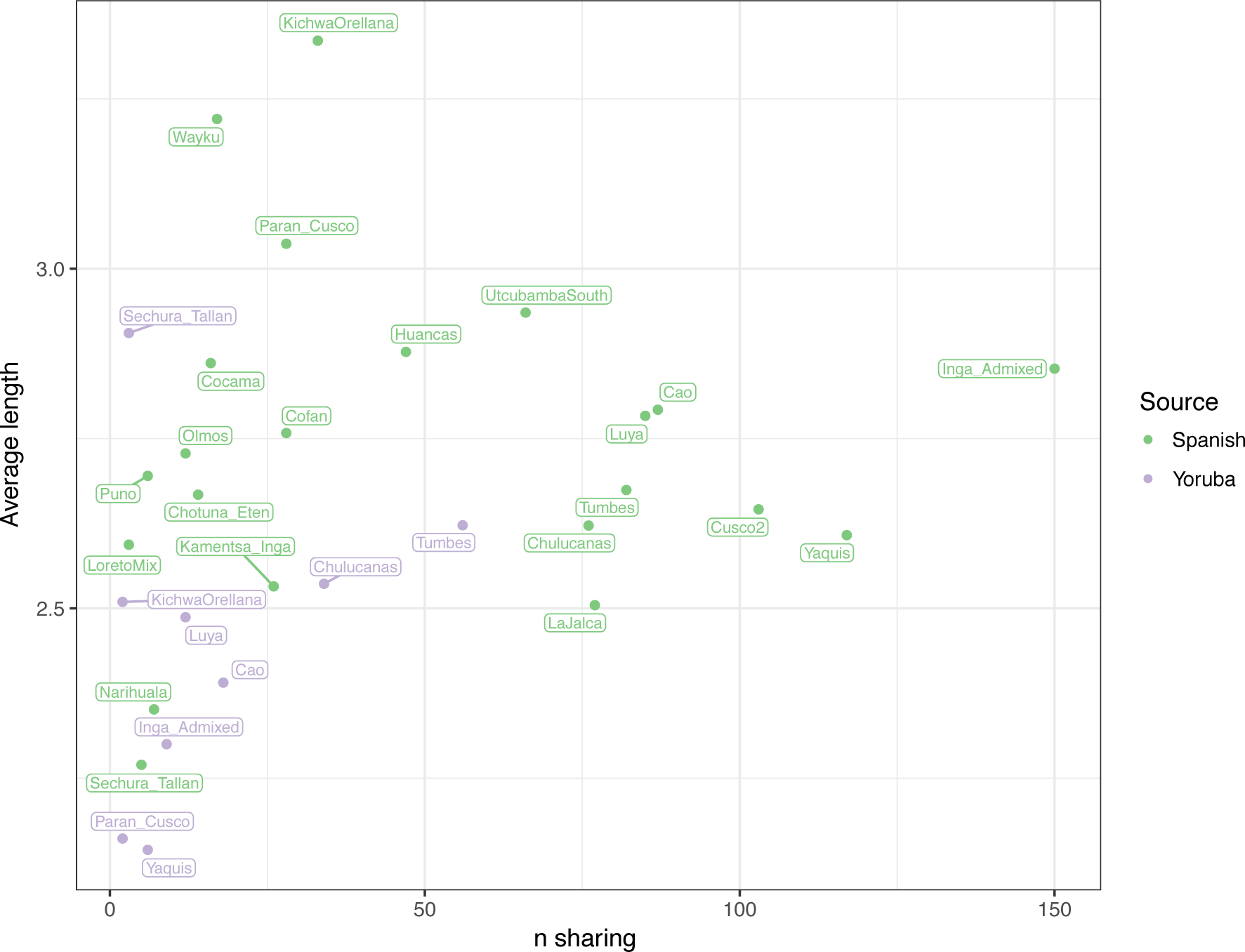
IBD block sharing with European and African sources. On the X axis, number of sharing events, and on the Y axis, average block length (in cM) per population.

